# SARS-CoV-2 methyltransferase nsp10-16 in complex with natural and drug-like purine analogs for guiding structure-based drug discovery

**DOI:** 10.1101/2024.03.13.583470

**Authors:** Viviane Kremling, Sven Falke, Yaiza Fernández-García, Christiane Ehrt, Antonia Kiene, Bjarne Klopprogge, Emilie Scheer, Fabian Barthels, Philipp Middendorf, Sebastian Kühn, Stephan Günther, Matthias Rarey, Henry N. Chapman, Dominik Oberthür, Janina Sprenger

## Abstract

Non-structural protein 10 (nsp10) and non-structural protein 16 (nsp16) are part of the RNA synthesis complex, which is crucial for the replication of severe acute respiratory syndrome coronavirus 2 (SARS-CoV-2). Nsp16 exhibits 2’-*O*-methyltransferase activity during viral messenger RNA capping and is active in a heterodimeric complex with enzymatically inactive nsp10. It has been shown that inactivation of the nsp10-16 protein complex interferes severely with viral replication, making it a highly promising drug target. As information on ligands binding to the nsp10-16 complex (nsp10-16) is still scarce, we screened the active site for potential binding of drug-like and fragment-like compounds using X-ray crystallography. The screened set of 234 compounds consists of derivatives of the natural substrate *S*-adenosyl methionine (SAM) and adenine derivatives, of which some have been described previously as methyltransferase inhibitors and nsp16 binders. A docking study guided the selection of many of these compounds. Here we report structures of binders to the SAM site of nsp10-16 and for two of them, toyocamycin and sangivamycin, we present additional crystal structures in the presence of a second substrate, Cap0-analog/Cap0-RNA. The identified hits were tested for binding to nsp10-16 in solution and antiviral activity in cell culture. Our data provide important structural information on various molecules that bind to the SAM substrate site which can be used as novel starting points for selective methyltransferase inhibitor designs.

## Introduction

Severe acute respiratory syndrome coronavirus 2 (SARS-CoV-2) can cause a mild to severe and life-threatening illness, termed coronavirus disease 2019 (COVID-19). It is similar to the disease caused by SARS-CoV and middle east respiratory syndrome coronavirus (MERS-CoV). Due to the global spread of SARS-CoV-2 and its impact on public health a call to action for a coordinated response worldwide was required. The World Health Organization (WHO) declared a pandemic in March 2020 (Zhou et al. 2020). Over 7 million cumulative deaths have been confirmed up to the end of 2023 (WHO 2023). Effective vaccines that reduced the number of critically ill patients needing hospitalization (Sandmann et al. 2021) were developed fast, however, immune escape variants have emerged (Dong et al. 2020, Parums 2023). Continuous efforts are in place to adapt these vaccines to address emerging variants (Grant et al. 2023). Approved drugs for COVID-19 treatment are available as well. However, they are mainly in use to treat patients at high risk of adverse outcomes due to the high costs, possible side effects and the potential emergence of drug-resistant strains (Pachetti et al. 2020; Food and Administration (FDA) 2021; 2021; Parums 2022; Bernal et al. 2022; Chatterjee et al. 2023; Parums 2023). Drugs directed toward highly conserved viral proteins are more likely to act on variants of concern unresponsive to vaccine-induced protection and are less likely to provoke the genesis of treatment escape variants (Pachetti et al. 2020; Chatterjee et al. 2023; Parums 2023).

SARS-CoV-2 is an enveloped β-coronavirus with a large (30 kb), complex, positive-sense single-stranded RNA genome. The genomic RNA contains a 5’-cap and a 3’-poly-A tail similar to host mRNAs, thereby preventing immune system activation (Menachery, Debbink, and Baric 2014; Daffis et al. 2010; Chang and Chen 2021; Bradrick 2017), enabling recognition by the host translation machinery, and increasing stability (Zhu et al. 2020; Furuichi and Shatkin 2000; Schwer, Mao, and Shuman 1998; Cougot et al. 2004; Lewis and Izaurflde 1997). The 5’ capping process involves several viral non-structural proteins (nsp) such as nsp13, RNA triphosphatase (TPase) helicase (Bouvet et al. 2010), the RNA guanylyltransferase (GTase) nsp12 (Walker et al. 2021; Yan et al. 2021), the *N*7-methyltransferase (*N*7-MTase) which is a complex of nsp10 and nsp14, and the 2’-*O*-methyltransferase (2’-*O*-MTase) complex nsp10-16 (Bouvet et al. 2010; Chen and Guo 2016; Ramanathan, Robb, and Chan 2016). The final steps of capping are the methylation of the 5’guanosine to form 7*N*-methyl guanosine (^m7^G) by nsp10-14 forming Cap0 and the methylation of the following nucleotide, often adenosine, at the 2’-*O* position to form Cap1 (Wilamowski et al. 2021). Both methyltransferases use the substrate *S*-adenosyl methionine (SAM) as a methyl donor and form a functional complex with the enzymatically inactive nsp10 (Decroly 2011; Gupta 2021; Minasov 2020). They are part of the viral replication and transcription complex (RTC) consisting of enzymes responsible for RNA replication (nsp12), processing enzymes (nsp14, nsp13, nsp16) and co-factors (nsp7, nsp8, nsp10) (Malone et al. 2022).

Early on, nsp10-16 has been identified as a potential drug target for SARS-CoV (Decroly et al. 2008; Chen et al. 2009; Byszewska et al. 2014) and has emerged as a therapeutic target against SARS-CoV-2 (Chang and Chen 2021; Liu et al. 2010; Dong, Zhang, and Shi 2008). It belongs to the class I methyltransferases and contains a Rossmann-fold, typical for methyltransferases, to bind SAM as the methyl donor and ^m7^GpppN-RNA (Cap0-RNA) as the methyl acceptor (Wilamowski et al. 2021). The large active site of nsp16 can be divided into two main parts, the SAM and the Cap0 site (figure 1) which previously have been termed the low-affinity binding site (LBS) and high-affinity binding site (HBS), respectively (Rosas-Lemus et al. 2020). In the context of this study, we further divide the SAM site into an adenosyl and a methionyl cavity (figure 1B). Several co-crystal structures of nsp10-16 with active site binders are available with ligands always binding into the SAM site and occupying both cavities (Krafcikova et al. 2020; Rosas-Lemus et al. 2020; Bobileva et al. 2021; Klima et al. 2022). Unlike other compounds, the previously reported nsp16 inhibitor SS148 extends further at position *N*7 of the heterocycle beyond the SAM site, here referred to as the extended *N*7 pocket (figure 1B).

**Figure 1.**
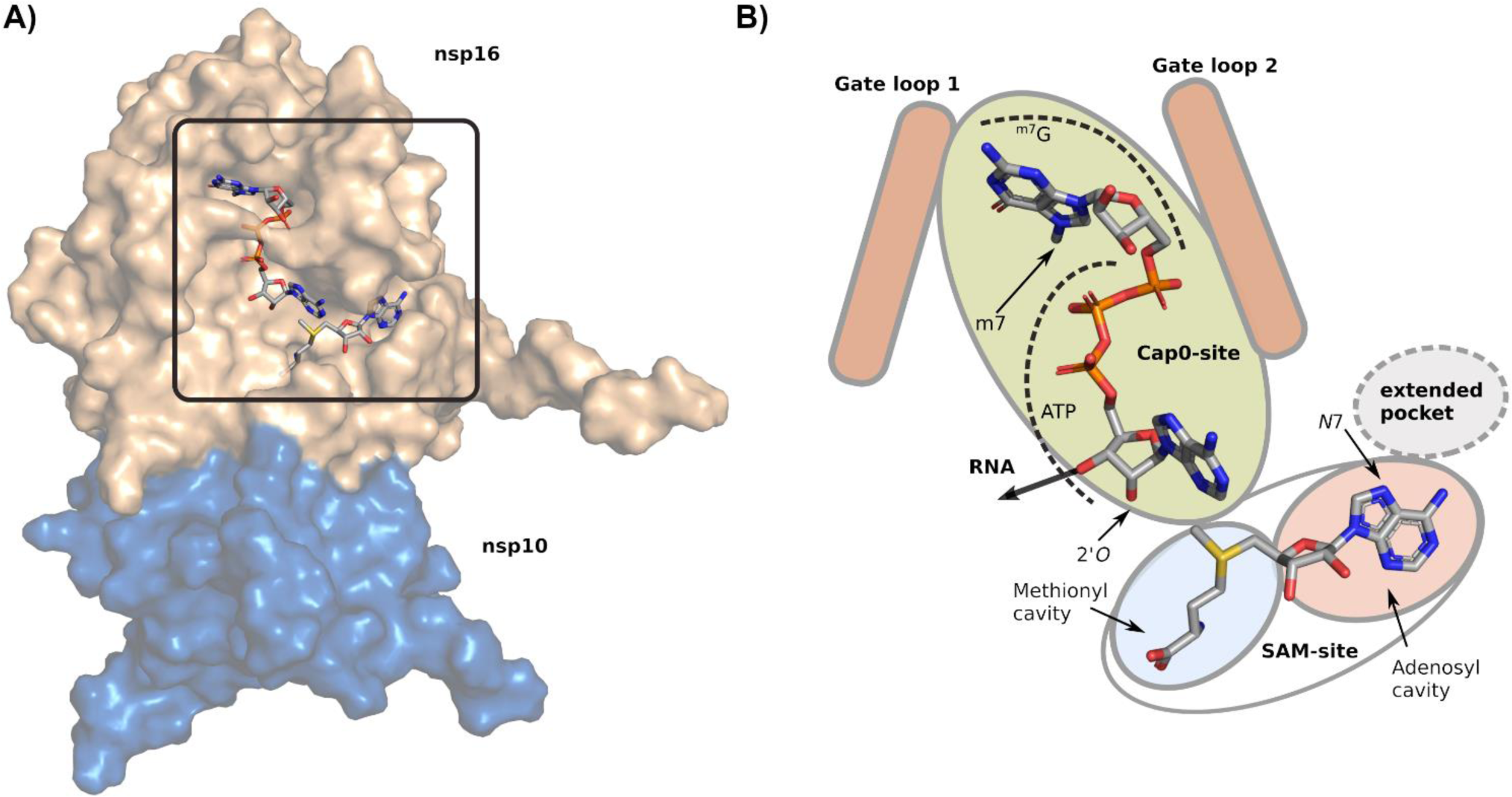
Nsp10-16 overview and active site architecture. Structural representation of nsp10-16 with nsp10 as blue and nsp16 as beige surface (based on PDB entry: 7JIB). A) The active site is highlighted with a black frame and shows the substrates SAM and Cap0-analog as stick models with carbon in gray, oxygen in red, sulfur in yellow, nitrogen in blue, and phosphate in orange. B) Schematic representation of the active site with ligands as stick models as in A). The relative positions of the gate loops and the sub division of the Cap0 and SAM site is shown as described in the text. The extended cavity at the SAM site extends from *N*7 at the adenosyl cavity.

The second substrate binding site, here termed the Cap0 site, accommodates the Cap0 molecule of the adjunct viral RNA. The Cap0 site can also bind a ^m7^GpppN molecule which includes only the first nucleotide, preferably adenosine (^m7^GpppA, Cap-analog). The Cap0 site is bracketed by two gate loops, i.e. gate loop 1 (Met6818 – Ile6838) and gate loop 2 (Met6929 – Asn6941) which are displaced upon binding of Cap0-analog or Cap0-RNA (Viswanathan et al. 2020). With SAM and Cap0 bound to the active site, the SAM methyl group and 2’-OH group of the Cap0 adenosine are in close proximity allowing for methyl transfer (Viswanathan et al. 2020). Additional ribonucleotides following the Cap0-adenosine increase the reaction efficiency, presumably because they induce a slight shift of the acceptor base resulting in a better position for the transfer reaction (Minasov et al. 2021).

Few nsp10-16 inhibitors are known to date. Sinefungin is an analog of *S*-adenosyl-L-homocysteine (SAH), the by-product of the methyl transfer reaction. In biochemical assays sinefungin has been shown to inhibit nsp10-16 (Otava et al. 2021; Devkota et al. 2021; Yazdi et al. 2021; Perveen et al. 2021) but it has a low inhibitory effect in cell culture-based assays (Bobileva et al. 2021; Bergant et al. 2022). Poor membrane permeability due to its zwitterionic nature has been suggested to cause this low activity (Ferreira de Freitas, Ivanochko, and Schapira 2019; Bobileva et al. 2021). In another study, the methionyl moiety of SAM was systematically exchanged and several compounds, mainly aromatic groups at this position, were identified as more potent than sinefungin but lacked selectivity (Bobileva et al. 2021). Recently, tubercidin has been suggested as a nsp16 inhibitor (Bergant et al. 2022). It is known to exhibit broad antiviral, antitrypanosomal, and antifungal effects but it is cytotoxic and lacks selectivity (Bergstrom et al. 1984; De Clercq et al. 1986; Eyer et al. 2016). Klima et al. 2022 identified the adenosine and tubercidin derivatives WZ16 and SS148, respectively (figure 2), as nsp16 inhibitors with an IC_50_ in the low micro molar range and crystal structures have been solved. These tubercidin-derived compounds are less cytotoxic and decrease the activity of nsp16 in biochemical assays (Klima et al. 2022; Schultz et al. 2022; Li et al. 2023) and in the case of SS148 also the activity of nsp14 (Devkota et al. 2021). Despite the knowledge of binders and inhibitors together with diverse *in silico* drug screens (Xu et al. 2020; Maurya et al. 2020; El Hassab et al. 2021; Vijayan et al. 2020) and an NMR fragment screen (Berg et al. 2022), no selective and effective nsp16 inhibitor has been identified to this point. Crystal structures of nsp10-16 from SARS-CoV, SARS-CoV-2, and MERS-CoV are available (Wilamowski et al. 2021; Chen et al. 2011, e.g. PDB entry 5YN6) and significantly contribute to the understanding of substrate binding and conformational changes upon Cap0 binding, catalysis and product release. The number of co-crystal structures with small molecule inhibitors is, however, limited and no X-ray compound screening has been undertaken unlike for other SARS-CoV-2 drug targets such as nsp14, PL_pro_, M_pro_, RdRp, and S glycoprotein (Günther et al. 2021; Manandhar et al. 2021; Kuzikov et al. 2021; Khalifa et al. 2021; Imprachim, Yosaatmadja, and Newman 2023).

**Figure 2.**
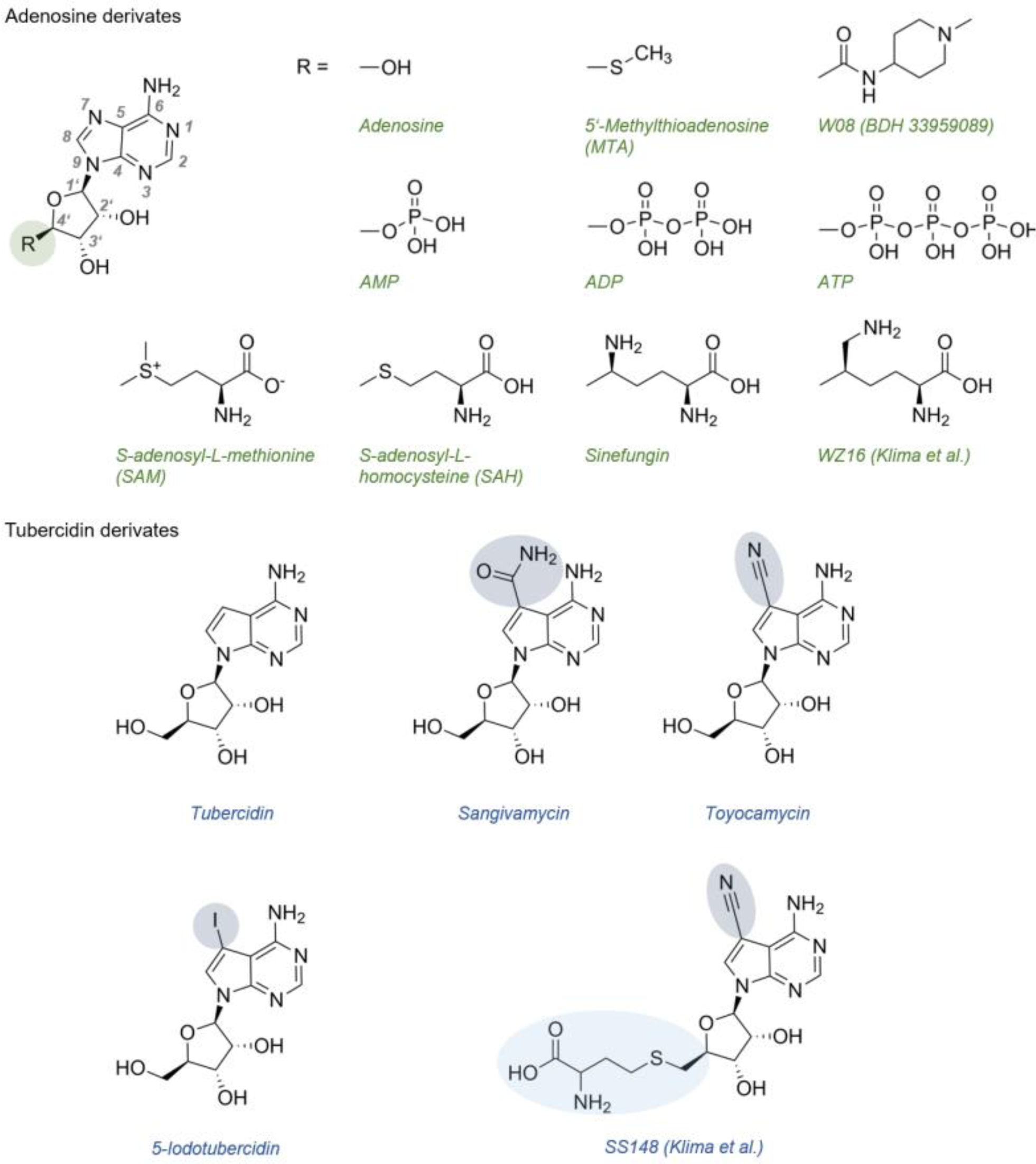
Chemical structures of SAM-site binders. The compounds on top represent adenosine (green labels) and on the bottom tubercidin derivatives (blue labels). Structures of SARS-CoV-2 nsp10-16 with SAM, SAH, Sinefungin, WZ16 and SS148 have been described previously. The protein complex structures of the remaining compounds are reported in this study. Chemical differences of the derivatives compared to tubercidin are highlighted with green/blue circles or ellipsoids.

Here, we explore the potential binding space of the nsp16 active site using a dedicated crystallographic screen of 234 compounds with derivatives of the natural substrate SAM as well as adenine analogues including previously described methyltransferase inhibitors. The selection of about half of these compounds was guided by molecular docking and the selection of a diverse subset by a fingerprint-based 2D similarity analysis of a large compound library targeting the SAM site. Further, we elucidate the structure of nsp10-16 in complex with tubercidin, which has been suggested as a potential nsp16 binder. We probed the binding of three additional tubercidin derivatives to nsp10-16 – namely toyocamycin, sangivamycin, and 5-iodotubercidin. Among the identified ligands binding to the SAM site, we discovered the adenosine analogue W08. Out of the *in silico* selected compounds for screening, W08 is structurally one of the most similar compounds compared to SAM, however, it binds unexpectedly in an ‘inverted manner’ compared to other analogs and extends into the extended *N*7 binding pocket. We explored the binding and antiviral effect of tubercidin and its derivatives which all bind to the nsp16 active site and whose *N*7 modifications extend into the extended *N7* pocket as well, retaining the conserved binding mode despite their modifications. Additionally, we obtained structures of toyocamycin and sangivamycin together with Cap0-analog and Cap0-RNA. It has been unclear if the entire SAM site needs to be occupied for the second substrate to bind. Toyocamycin and sangivamycin are lacking the methionyl moiety of SAM. The toyocamycin/sangivamycin Cap0-analog/-RNA structures show that the Cap0-binding site can be occupied independently of the methionyl moiety. Our results provide important information required to understand the structural variety of compounds that are able to occupy the nsp16 active site which will enable the design of more specific antiviral methyltransferase inhibitors in the future.

## Results

### SAM is replaced by derivatives of SAM, adenosine, and tubercidin

Crystals of nsp10-16 grew within one to two weeks to 75-150 µm size under crystallization conditions reported by Rosas-Lemus et al. 2020 and crystal structures with ligands were refined against X-ray diffraction data at resolutions between 1.6 to 2.2 Å (table S 1). In total, 234 compounds were screened including two libraries from ASINEX, selected SAM fragments, tubercidin derivatives and known nsp16 inhibitors (figure S1) and identified 10 hits in the SAM site. Additionally, we identified several hits in the Cap0 site which will be described in a future manuscript (table S 2). All structures show the same overall fold as previously described (Viswanathan et al. 2020; Rosas-Lemus et al. 2020). In the absence of other bound compounds, clear SAM electron density is observed in the active site although no SAM was added. The capturing of SAM during protein expression in *E.coli* has been observed previously (Lin et al. 2020). The soluble fraction of nsp10-16 contains about 60 % bound SAM, as confirmed by isothermal titration calorimetry (ITC) (figure S 2A). However, the high occupancy in the crystal (modeled as 100 %) suggests that only the SAM-bound form crystallized. Attempts to remove SAM from soluble nsp10-16 prior to crystallization by extensive dialysis using charcoal led to 80 % ligand free-enzyme (compared to 40 % without dialysis) as shown by ITC (figure S 2B). However, crystallization trials with this SAM-reduced nsp10-16 did not yield crystals even when supplemented with SAM. The compounds identified as nsp16 binders in the crystal screen were soaked into crystals containing SAM and partially or fully replaced SAM. Partial replacement was observed for the ligands adenosine triphosphate (ATP), adenosine diphosphate (ADP), and W08 (ASINEX ID BDH 33959089). The combined occupancy of SAM and the respective ligand sums up to 100 %, suggesting that all active sites of the enzyme in the crystal are occupied. The other identified SAM site binders fully replace SAM. The binding of all tested compounds induces slight changes in the backbone root-mean-square deviation of atomic positions (RMSD) with the highest flexibility in the gate loops (figure S 3).

### SAM site binders occupy the adenosyl cavity and bind to an extended pocket

Adenosine, 5’-methylthioadenosine (MTA), and tubercidin are structurally similar (figure 2) and electron density is observed in the adenosine cavity while SAM is fully replaced (figure S 4). The active site interactions are identical to those of the adenosyl moiety of SAM where the following protein and ligand atoms are within hydrogen-bonding (H-bonding) distances: Asp6912 carboxyl oxygen and exocyclic nitrogen at *C*6; Cys6913 backbone nitrogen and *N*1; the carboxyl oxygens of Asp6897 and both ribose hydroxyl groups; Tyr6930 backbone nitrogen with cyclic oxygen of the ribose (figure 3). Additionally, the backbone carbonyl oxygen of Tyr6930 is within H-bonding distance to the 5’-OH of tubercidin, adenosine and MTA. These hydroxyl groups show alternative conformations modeled with almost equal occupancies either in the ‘up’ (in close proximity to Tyr6930, occupancy: 40 %) or in the ‘down’ position (occupancy: 60 %) with the oxygen at the same position as the sulfur atom in SAM (figure 3).

**Figure 3.**
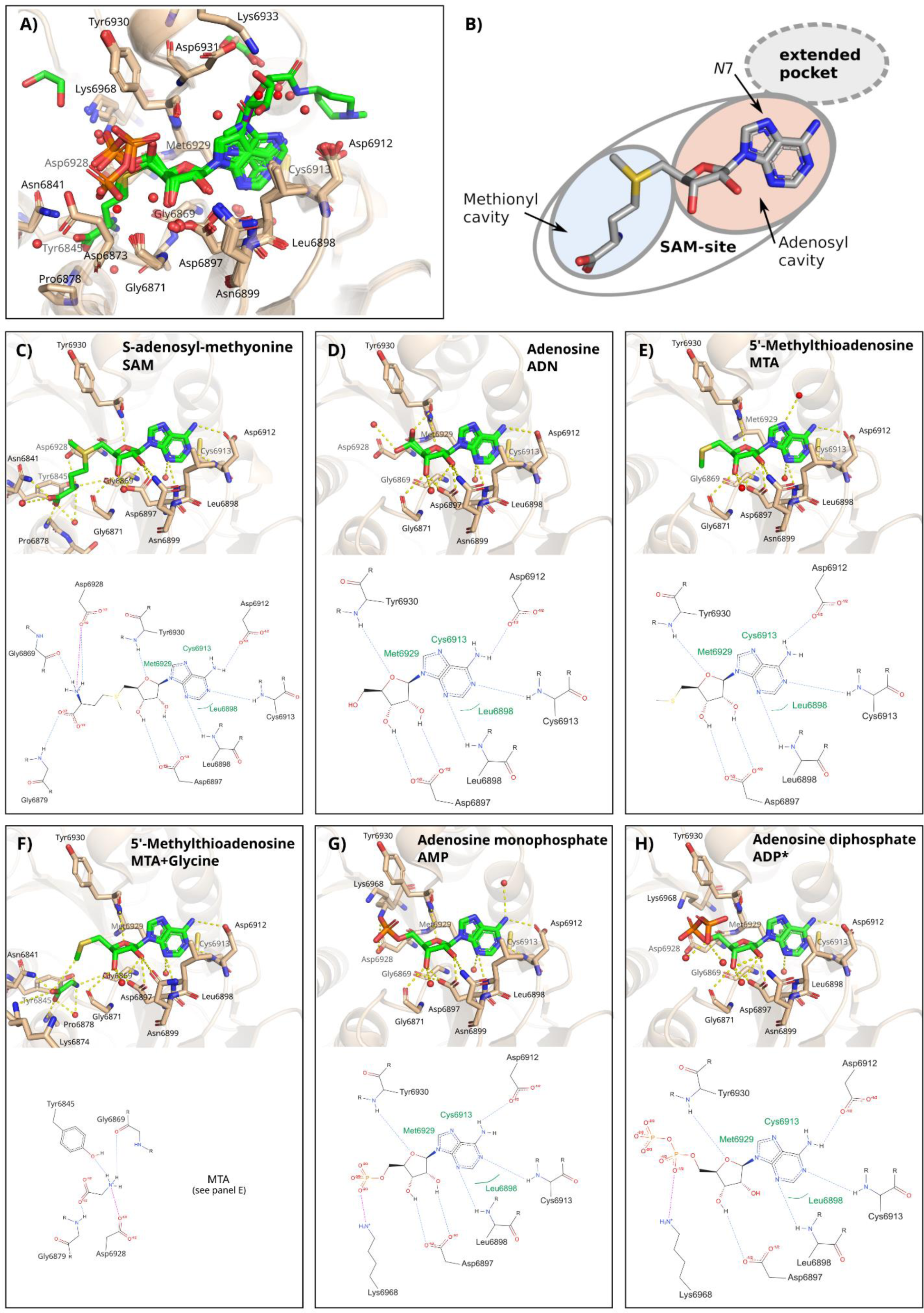
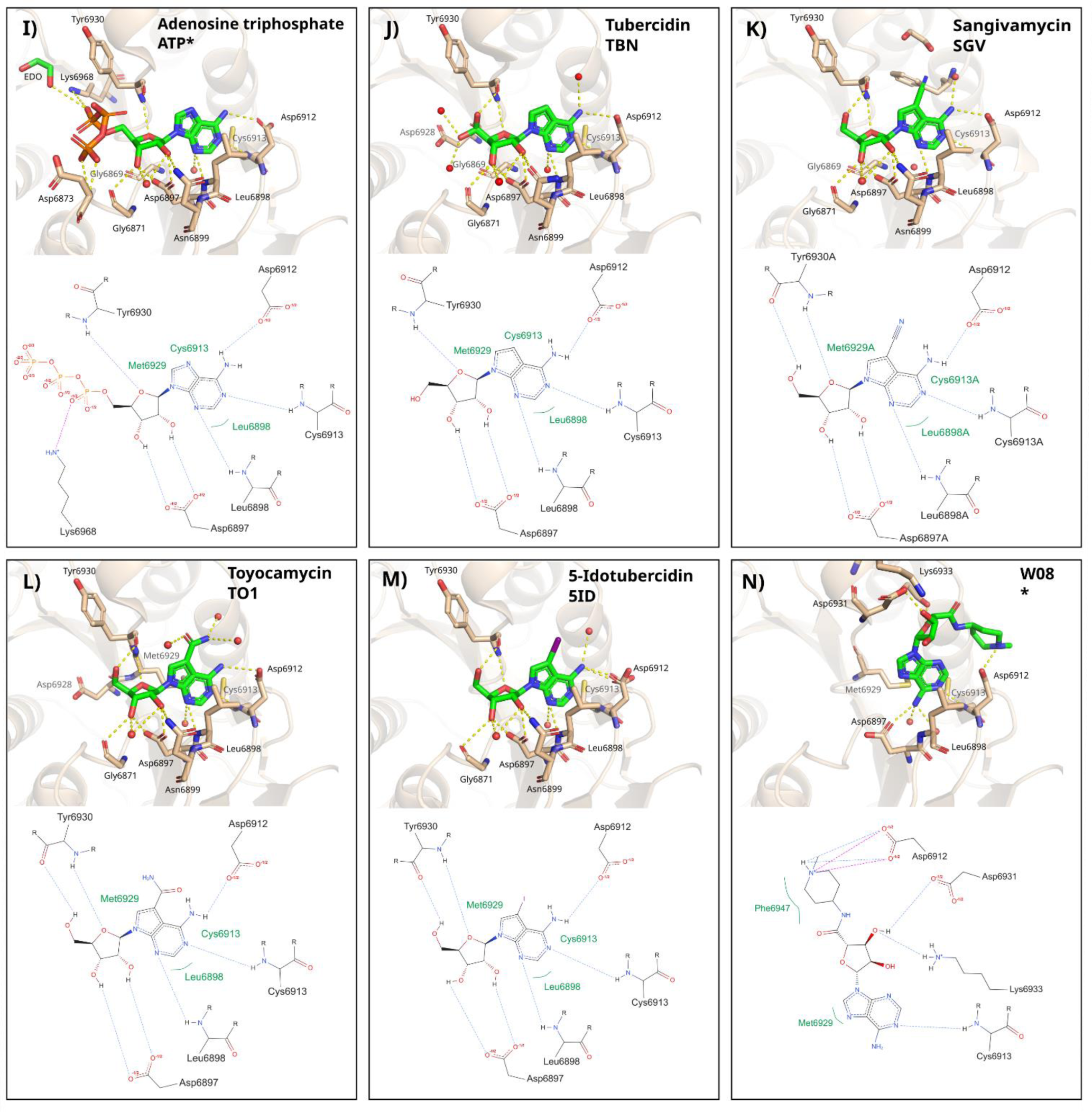
Overview of interactions of ligands with nsp10-16. A) Overlay of ligands (stick representation) bound to SAM binding site of nsp16 (cartoon representation). B) SAM site architecture adapted from figure 1 B). C) - M) interaction of different ligands with nsp16. The top of each panel shows the ligand poses in the crystal including possible interactions as yellow dotted lines. On the bottom, possible interactions as 2D-plots only including relevant residues without the protein backbone are shown. Plots were created and edited with PoseView and PoseEdit (https://proteins.plus, Schöning-Stierand et al. 2022; Diedrich et al. 2023). Complex structures with partial occupancy by SAM are marked with an asterisk. For clarity, SAM is not shown even if present in these crystal structure.

Furthermore, Met6929 might be involved in hydrophobic contacts with *C*2 of the purine of tubercidin and adenosine. In structures including SAM, this residue is in close proximity to the SAM sulfur atom. One to two water molecules are near to the exocyclic nitrogen of tubercidin in the ‘down’ conformation and might be involved in additional H-bonding. The binding mode of tubercidin agrees with the proposed *in silico* binding pose by Bergant et al. 2022.

To understand whether the binding of SAM can be imitated by two individual smaller fragments, we sequentially soaked MTA and glycine, resulting in ligand electron density for both compounds. In this structure, the orientation and interactions of MTA with the active site residues are identical to the structure of MTA alone (figure 3E, F). Glycine binds to the methionyl cavity which is occupied by an ethylene glycol in the structure with MTA alone. The potential interactions of the bound glycine with the protein are the same as the glycine corresponding part of the methionyl moiety of SAM. Hence the presence of glycine in the active site is most likely, showing that SAM binding can be imitated by the fragments MTA and glycine.

The tubercidin derivates toyocamycin, sangivamycin and 5-iodotubercidin share the same orientation as tubercidin and the above-mentioned possible interactions with surrounding residues and waters are preserved (figure 3 J-M). None of the 5’substituents of these derivatives have any surrounding interacting residues except for the amide group of sangivamycin that is in H-bonding distance of an ethylene glycol and a water molecule (figure 3K) or three water molecules in the case of the nitrile group of toyocamycin (figure 3L). The electron densities of the 5’-OH groups of 5-iodotubercidin and sangivamycin are interpreted as in the ‘up’ position but the ‘down’ position cannot be excluded (figure 3K, M). Vice versa, toyocamycin was modeled in ‘down’ position although the ‘up’ position cannot be excluded (figure 3L). The positions of tubercidin derivatives are very similar except for the aromatic heterocyclic ring of sangivamycin that is slightly shifted, resulting in a 0.4 Å offset of the 5’-C compared to tubercidin. The neighboring residue Asp6931 has also shifted away by 0.2 Å in this structure compared to tubercidin. In summary, a strong effect of 5’substituents on active site residues could not be observed. This coincides with the observations for the binding and protein-ligand interaction of W08.

W08 is an adenosine analog and was on a low rank in the docking studies but is the third most similar compound to SAM in the library of docked compounds from ASINEX. In the crystal structure, electron density is present for W08 (occupancy: 57 %) and SAM (occupancy: 43 %). Surprisingly, the binding pose of W08 shows a 180° rotation compared to the SAM scaffold. As a consequence of the rotation, the ribose moiety points out into the extended *N*7 pocket towards the same position as the *N*7 substituents of the tubercidin derivates (figure 3N). The contacts between the adenosine moiety of W08 and the active site residues differ from those observed with SAM (figure 3A, N). The exocyclic nitrogen of W08 is in close proximity to the carbonyl side chain atoms of Asp6897, whereas in SAM complexes the carboxyl moiety is in contact with the nitrogen of Asp6912. The cyclic *N*1 is, similar to SAM, in H-bonding distance to the backbone nitrogen atom of Cys6913. In the extended pocket, occupied by the piperidinamide ribose of W08, the 4’-OH group of the ribose is in H-bonding distance to the carbonyl oxygens of Asp6931 and Lys6933; the cyclic amine of the substituent is close to a carboxyl oxygen of Asp6912 and Phe6947. The latter might be involved in hydrophobic contacts with the piperidine ring (figure 3N).

The adenosine phosphates AMP, ADP and ATP were included in the screen to probe the Cap0 binding site as these are natural ‘fragments’ of Cap0 (figure 2B). Ligand electron density was, however, only observed in the SAM-binding site (figure S 4E-G). Their adenine moiety aligns with the equivalent substructure of SAM (figure 3A). The phosphates of AMP, ADP, and ATP are flexible and extend out of the SAM pocket and do not occupy the methionine cavity (figure 3G-I). In all cases, Lys6968 is in close proximity to the α-phosphate oxygen and the γ-phosphate oxygens of the compounds that might also interact with the backbone nitrogen of Asp6873 (figure S 4E-G). Partial SAM density is still observed besides ADP (occupancy: 59 %) and ATP (occupancy: 58 %).

### In silico compound selection

For eight compounds with the highest ECFP4-based 2D similarity to SAM in the ASINEX library, electron density was not found in the SAM site after soaking experiments. Only W08 was identified as a SAM site binder and was selected based on its high 2D similarity to SAM is W08 which, however, binds in an unexpected orientation regarding the common molecular substructure of W08 and SAM.

Furthermore, no hits in the SAM site were found for the 93 compounds selected based on the molecular docking. For two compounds that include a theophylline substructure, partial electron density was found at the ^m7^G-position in the Cap0-site (results not shown). The docking with JAMDA (Flachsenberg et al. 2023) led to compounds with higher scores than the one observed for SAM. However, the compounds are much larger than known binders and extend into the methionyl cavity or Cap0 binding site. Examples of poses are provided in figure S 5. We applied the crystal channel analysis tool LifeSoaks (Pletzer-Zelgert et al. 2023) to see whether soaking of the proposed compounds might be hindered by narrow channels in the crystal. However, an analysis of PDB entry 8BSD shows that a broad channel with a 48.9 Å radius leads through the crystal and the binding site is freely accessible with an outer active site radius of 15.7 Å. The main differences between the selected compounds and known binders is the missing aliphatic character of the moiety attached to the ribosyl moiety of the compounds. Additionally, several compounds harbor other bicycle aromatic moieties instead of the adenine in SAM. Recent studies suggest that hydrophilic compounds are challenging regarding pose prediction with JAMDA (Flachsenberg et al. 2023). This finding might explain why molecular docking did not lead to the identification of novel binders for nsp10-16.

### Replacement of SAM with tubercidin derivatives still allows binding of the Cap0- substrate

Compared to the native substrate SAM, tubercidin and its tested derivatives lack the methionyl moiety (figure 2). It is unclear if that part of the ligand in the SAM site affects the binding of the Cap0-RNA. To test if the second substrate, Cap0, can bind in the presence of selected SAM-site binders, toyocamycin and sangivamycin were soaked sequentially with Cap0-analog or Cap0-RNA (^m7^GpppA UUAAA GGUUU AUACC UUCCC AGGUA) into nsp10-16 crystals. The resulting electron densities were compared to structures with SAM and Cap0-analog/Cap0-RNA. A structure containing SAM and Cap0-analog has been published previously (PDB entry 7KOA, Wilamowski et al. 2021) but from crystals grown under different conditions. For better comparability with structures from our study, we obtained crystals containing the two substrates under the same conditions as ligand-soaked crystals. Here we found that ethylendiamine tetraacetic acid (EDTA) that had been used to slow down the methyl transfer reaction can also bind to the surface of nsp10-16 (figure S 6). More information on this finding can be found in the supplementary material. Nevertheless, the shift of the gate loops as seen in the substrate structures with SAM and Cap0-analog present (Figure 4 A) is observed when SAM is replaced by either of the tubercidin derivates, toyocamycin or sangivamycin (Figure 4B). In all these structures the binding of the Cap0-analog and Cap0-RNA causes a shift of 1.5 Å in gate loop 1 for Tyr6828 (based on the backbone nitrogen position) and a shift of 2.4 Å of the Cα from Lys6935 and 2.1 Å of Cα from Pro6932 in gate loop 2 (figure 4A, B; S 2). There are no obvious differences in structures or interactions visible in the crystal structure from the tubercidin derivatives plus Cap0-analog/Cap0-RNA and the reference structure with SAM and Cap0-analog (figure 4A, C, D), suggesting that the binding of smaller compounds that do not occupy the methionine cavity also allows the native motions of the gate loops.

**Figure 4.**
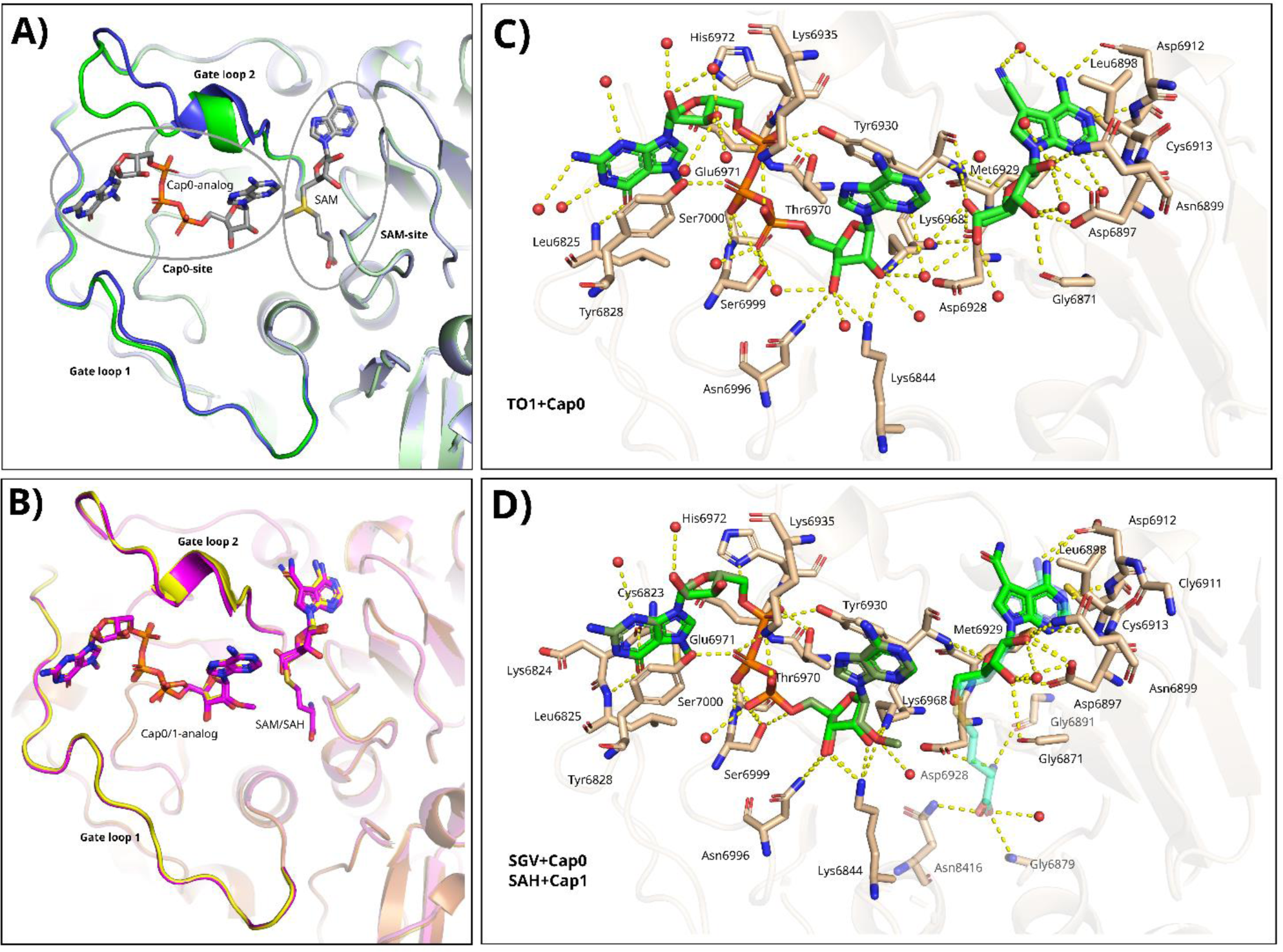
Cap binding and gate loop motions. A) Gate loop conformations for nsp16 with bound SAM (blue ribbon) in overlay with the SAM and Cap0-analog bound structure (green ribbon). SAM and Cap0-site positions are indicated by circles. B) Overlay of nsp10-16 structures with bound toyocamycin and Cap0-analog (yellow ribbon and sticks) with the structure resulting from soaking sangivamycin and Cap0-analog (pink ribbon and sticks). In the latter structure, SAM and SAH are present aside from sangivamycin and a fraction of the Cap0- analog was converted into Cap1. The orientation is identical to A). C) Interactions of Cap0-analog and toyocamycin (TO1) with nsp16. D) Interactions of SAH/sangivamycin (SGV) and Cap0/Cap1-analog with nsp16.

For toyocamycin together with Cap0-analog and Cap0-RNA, clear density is present (figure S 3M, N). Toyocamycin replaced SAM, although a remaining fraction of SAM cannot be excluded. Density in the allosteric binding site in the structure with Cap0-analog was interpreted as 7-methyl-guanosine-5’-triphosphate (MGP) as in other nsp10-16 crystal structures (PDB-entries 7JIB, 7JHE, 6WRZ, 6WVN). The crystal structure with toyocamycin and Cap0-RNA contains a Mg^2+^ ion in the proximity of Cap0. In the case of sangivamycin soaked with the Cap0-analog and Cap0-RNA, the resulting densities are more ambiguous. The structure with the analog contains an overlay of Cap0-analog (occupancy: 46 %) and the product m7GpppAm (Cap1-analog) (occupancy: 54 %) and in the SAM site an overlay of sangivamycin (occupancy: 36 %) with SAH (occupancy: 64 %). In the structure with the Cap0-RNA, the RNA molecule was modeled with 100 % occupancy and in the SAM site sangivamycin (occupancy of 39 %) is present together with SAM (occupancy: 61 %).

### Tubercidin and its derivatives reduce viral titers

All identified binders of nsp16 (Adenosine, MTA, AMP, ADP, ATP, tubercidin, sangivamycin, toyocamycin, 5-iodotubercidin) were tested for their *in vitro* antiviral activity at 10 µM concentrations except for W08 since it was no longer available for purchase. SARS-CoV-2 infectious particles in the supernatant of treated infected Vero E6 cells were quantified via immunofocus assay while the effect of the compounds on cell viability was determined employing the cell counting kit-8 (CCK-8) method. Only tubercidin derivatives reduced viral titers by at least ten-fold, although at a clear expense of cell viability (figure S 7). Notably, sinefungin known as a pan-methyltransferase inhibitor and widely used as a control in nsp10-16 biochemical assays (Otava et al. 2021; Devkota et al. 2021; Yazdi et al. 2021; Perveen et al. 2021) did not decrease virus replication. A plausible explanation for the lack of virucidal properties of sinefungin is its poor cell permeability (Bobileva et al. 2021).

To calculate the selectivity index (SI) of tubercidin substituents, we estimated their half-maximal effective concentration (EC_50_) and half-maximal cytotoxic concentration (CC_50_) in dose-response experiments. All tubercidin modifications reduced viral titers in a dose dependent manner (figure 5). However, due to its ability to reduce infectious particles by more than a hundred-fold exhibiting a SI greater than five, under our experimental conditions, we exclusively consider sangivamycin to have a selective effect on SARS-CoV-2 replication with an EC_50_ of 0.01 µM (table 1). Although tubercidin reduces viral titers by more than hundred-fold, it presents limited therapeutic value due to an unfavorable toxicity profile demonstrated in cell-based assays by a SI of 3.75 (figure 5, table 1). Meanwhile, the reduction of viral titers detected at 10 µM and higher concentrations of toyocamycin and 5-iodotubercidin are largely the effect of cytotoxicity (figure 5, **Table 1**).

**Figure 5.**
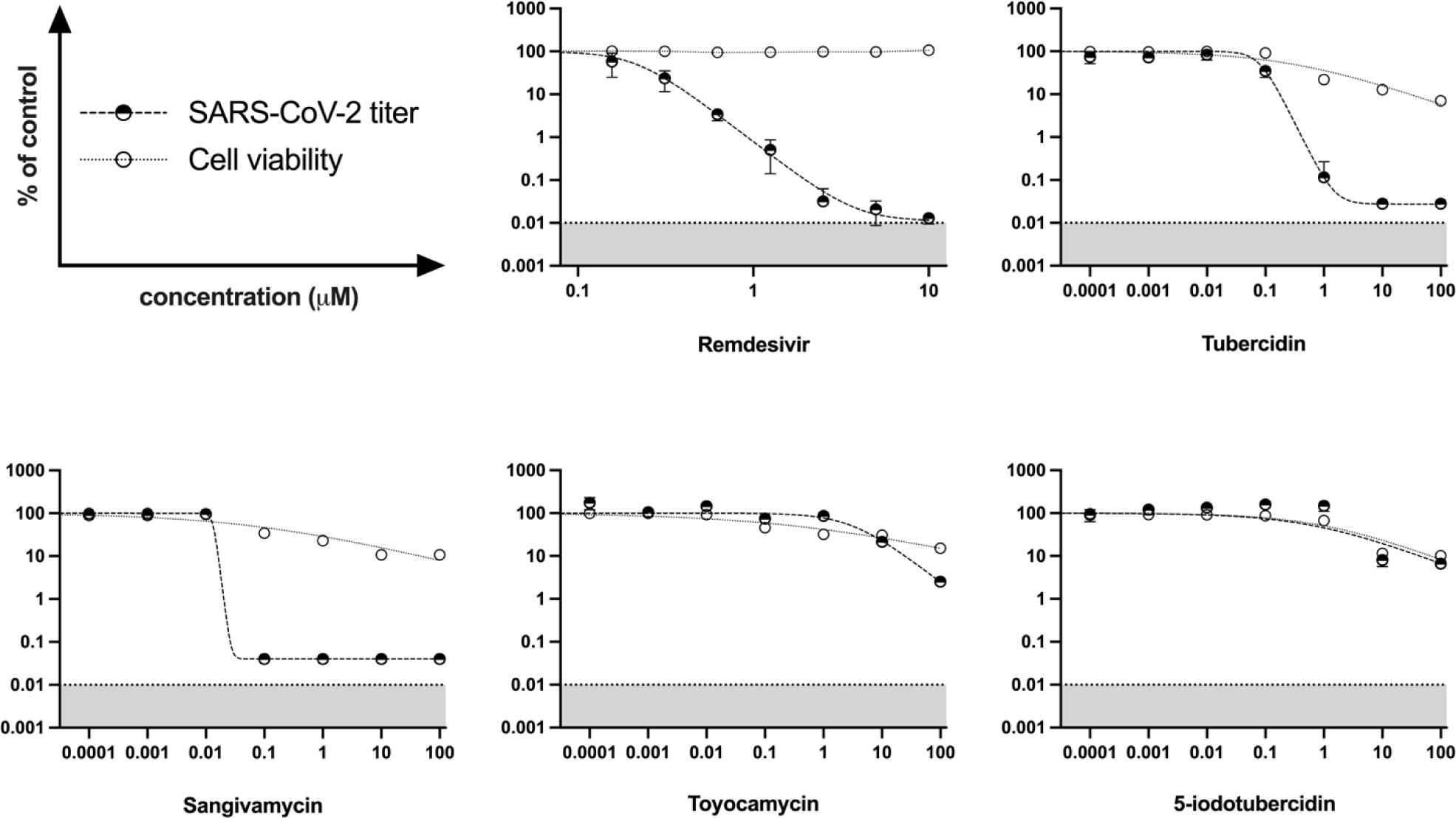
Effect of selected compounds on SARS-CoV-2 replication in Vero E6 cells. The viral titers (half-solid circles) and cell viability (empty circles) were determined by immunofocus assays and the CCK-8 method, respectively. Individual data points represent means ± SD from three independent replicates in one experiment. Values below the methodological detection limit are shaded in gray. The dose-response curves were generated by fitting the data to the sigmoidal four parameter logistic function with 1/Y^2^ weighting using GraphPad Prism version 9.5.1 for Mac OS X (GraphPad Software, San Diego, California USA, www.graphpad.com). The maximum values were fixed to 100 %, and the minimum values were constrained to be equal to or greater than the methodological detection limit. Remdesivir was used as positive control.

**Table 1.** Cytotoxicity, *in vitro* antiviral activity, and selectivity of selected compounds against SARS-CoV-2. CC_50_- half-maximal cytotoxic concentration; EC_50_- half-maximal effective concentration; SI- selectivity index. Cell viability and viral titers were determined by the CCK-8 method and immunofocus assays, respectively. Compounds that reduced infectious particles by at least a hundred-fold in combination with SI greater than five are considered antivirally active and are marked with an asterisk. Except for the SI, values were calculated from three independent replicates in one experiment by fitting the data to the sigmoidal four parameter logistic nonlinear regression function.

| Compound | Cytotoxicity<br>CC<sub>50</sub> [μM] | Antiviral activity<br>EC<sub>50</sub> [μM] | Selectivity index<br>SI (CC<sub>50</sub>/EC<sub>50</sub>) |
| --- | --- | --- | --- |
| Remdesivir * | >10 | 0.22 | >45.46 |
| Tubercidin | 0.3 | 0.08 | 3.75 |
| Sangivamycin * | 0.07 | 0.01 | 7 |
| Toyocamycin | 0.34 | - | - |
| 5-iodotubercidin | 1.07 | - | - |
| MTA | >100 | - | - |
| Sinefungin | >100 | - | - |

### Binding site comparison reveals structurally similar SAM pockets to nsp16 in human methyltransferases

Given the high cytotoxicity of the successfully soaked binders of nsp10-16, we set out to identify potential off-targets of the compounds. To this end, we applied SiteMine (Reim et al. 2024) to search for similar binding sites of SARS-CoV-2 nsp10-16 (figure S 8A) in structures of human methyltransferases. Figure S 8B shows the distribution of similarity scores obtained for searching in a methyltransferase-focused subset of the PDB (see Methods). The most significant similarities were found to SAM-complexed binding sites of the mitochondrial rRNA methyltransferase 2 (MRM2, UniProt accession: Q9UI43, PDB entry 2NYU; figure S 8C) and Cap-specific mRNA (nucleoside-2’- *O*-)-methyltransferase 1 (HMTR2, UniProt Accession: Q8N1G2, PDB entry 4N49; figure S 8D). MRM2 fulfills crucial functions in the mitochondrial machinery and mutations in the enzyme cause MELAS-like clinical syndrome (Garone et al. 2017). The function of HMTR2 is poorly understood (Werner et al. 2011). Besides these two hits, we found 30 further enzymes with similar predicted or ligand-based sites. Several are critical in the human organism, e.g., tRNA methyltransferases and ribosomal rRNA processing enzymes. We visually inspected all binding site matches with a score of at least 31 (139 matches that can be clustered into 32 distinct proteins). Details regarding this analysis can be found in the supplementary material. We further investigated the similarities and differences between the SAM binding site of nsp10-16 and the best-scored hits (figure S 8). Intriguingly, we found highly conserved residues involved in hydrogen bond interactions with SAM (figure S 8). However, the surface and physicochemical properties in the direction of the *N*7 extension vary considerably (figure S 8). This finding might provide new starting points for the design of SAM-based selective inhibitors.

### Ligands bind to the nsp16 active site with low affinity

To assess ligand affinities for the nsp16 active site, a diverse set of methods was employed including nano differential scanning fluorimetry (nanoDSF), microscale thermophoresis (MST), ITC, ITC displacement assay, and spectral shift (NanoTemper Technologies). In brief, nanoDSF measurements with 10 nM protein and 1 mM ligands showed an increased stability of nsp10-16 with bound Cap0-analog (figure S 9C). SAM and SAH did not induce a thermal shift. Tubercidin and its derivatives seem to destabilize the protein complex (figure S 9A, B, D). Neither ITC (figure S 10) nor any other method tested showed measurable binding of the tested ligands within the investigated ligand concentrations applied, except for the substrates SAM and Cap0-analog. Spectral shift is not suitable for measuring nsp10-16 compound binding since no binding of the substrates SAM and Cap0-analog could be detected. In an attempt to measure the presumably weak binding of sangivamycin, an ITC displacement experiment was conducted where 2 mM sinefungin (as a “strong” binder) was titrated into 42.5 µM protein that had been supplemented with 2 mM sangivamycin in advance. A control titration with sinefungin into protein was subtracted. The resulting K_D_ value of sangivamycin is > 1 mM (figure S 11).

Taken together, despite clear density visible in difference electron density maps for ligands binding to the nsp16 active site, we could not determine exact apparent K_D_ values which are, however, estimated to be in the milli molar range.

In an effort to establish an MST displacement binding assay with SAM like molecules with an attached fluorophore, we obtained a crystal structure of nsp10-16 with the fluorescent RNA methyltransferase probe 5-FAM-triazolyl-adenosyl-Dab (FTAD) (Zimmermann et al. 2022) (figure S 12). More detailed information on binding assays and the FTAD crystal structure can be found in the supplementary material.

## Discussion

The lack of knowledge about binding fragments and small molecules to SARS-CoV-2 nsp10-16 has restricted the search and design of new and specific inhibitors. This study shows an approach similar to fragment screening with small drug or drug-like molecules to find new potential leads that can be further developed into selective inhibitors. Our screening approach using X-ray crystallography focused on SAM analogs but also included other smaller adenosine analogs like tubercidin and its derivatives. Unexpectedly, the compound W08 from an ASINEX library binds in a reverse orientation compared to all other identified binders and occupies an extended pocket of the SAM binding site instead of the methionyl cavity although it is one of the three most similar compounds to SAM in the library. This extended pocket might be a promising starting point to modify existing or develop future inhibitors to gain higher affinity and selectivity. Similar to W08, the tubercidin derivatives sangivamycin, toyocamycin and 5-iodotubercidin expand into this extended pocket, as do the already known inhibitors WZ16 and SS148 (Klima et al. 2022).

Despite clear compound electron densities in the nsp16 active site, binding constants could not be measured for tubercidin and tubercidin derivatives in any biochemical assay used in this study, except for sangivamycin which has a K_D_ of > 1mM measured in an ITC displacement assay where sangivamycin was displaced by sinefungin. These compounds, however, showed different degrees of SARS-CoV-2 replicative attenuation in cell-based assays where all proved to be cytotoxic at a high nano molar range. Only the therapeutic efficacy zone of sangivamycin indicates a selective and potent activity against the virus. Tubercidin has been suggested as a nsp16 inhibitor. However, its antiviral effect has been characterized as a bispecific MTase inhibitor capable of reducing the enzymatic activity of the human cap methyltransferase 1 (HMTR2, UniProt: Q8N1G2) that can rescue SARS-CoV-2 mRNA methylation to a certain extent when the viral nsp16 catalytic function is impaired (Bergant et al. 2022). From the SiteMine binding pocket analysis HMTR2 has also been identified as one of the most similar enzymes to nsp16. Off-target effects on the viral site are very likely for 5-iodotubercidin and toyocamycin. 5-iodotubercidin has already been shown to target the multifunctional nsP1 that synthesizes Cap0 in the Chikungunya virus (Mudgal, Mahajan, and Tomar 2020), and it further inhibits SARS-CoV-2 RNA-dependent RNA polymerase (RdRp, nsp12). Toyocamycin can inhibit polymerases of other viruses (Bergstrom et al. 1984; Wang and Yang 2020) but has not been tested yet against the SARS-CoV-2 RdRp. Another tubercidin derivative, 5-hydroxymethyltubercidin, can inhibit viral replication potentially by blocking the viral RdRp and with that the RNA synthesis of coronaviruses, including SARS-CoV-2 (Uemura et al. 2021). Targeting several viral proteins might prove beneficial for treatment, however, off-target effects in the patient should be avoided. Although restricted to publicly available structures, our analysis points towards more off-targets in human proteins of nsp10-16 inhibitors and highlights differences to be exploited in upcoming design studies to potentially reduce toxicity. On the other hand, the knowledge of similar sites might also help to develop novel ideas for selective inhibitors (e.g., PDB entry 6CKC - protein arginine N-methyltransferase 5 in complex with the selective and potent inhibitor LLY-283).

Although tubercidin derivatives may not target nsp16 specifically, these compounds can be used as starting points for further modifications to gain potency, specificity, and selectivity which has been demonstrated, e.g. by Klima et al. (2022) with the tubercidin derivative SS148 that inhibits nsp16 activity (IC_50_ of 1.2 µM with 1.7 µM ^3^H-SAM and 0.8 µM biotinylated-Cap0-RNA substrate concentrations). SS148 can be described as a merging product of toyocamycin and the homocysteine moiety of SAH. The native substrate SAM and the product SAH are built from adenosine or a methionine and homocysteine moiety, respectively. We identified adenosine as a binder in the active site of nsp16 in the crystal. Although we could not determine the binding affinity, its density is clearly visible in the crystal structure and can be used as a minimal building block for drug design.

The inhibitors sinefungin and its derivate WZ16 identified by Klima et al. (2022) have detectable affinities and inhibitory activities in solution. They both occupy the methionyl cavity which might decrease compound affinity. Bobileva et al. (2021) systematically replaced the methionyl moiety of SAM, resulting in several potent inhibitors that contain aromatic structures instead of the methionine. Especially *m*-benzoic acid derivatives show promising results despite lacking selectivity. These chemical scaffolds are valuable building blocks. In line with these observations, it is known that fragments and fragment-like compounds typically show weak binding and inhibition with K_D_/K_I_ values in the micro molar to milli molar range (Maveyraud and Mourey 2020). Although exceeding the aim of this study, a fragment merging approach might result in potent new inhibitors where the occupation of the methionyl cavity together with the extended pocket increases affinity and selectivity.

An intriguing approach is to target both the SAM and Cap0-site of nsp16. Several compounds selected from the docking study are larger than the known binders and were predicted to extend from the SAM to the Cap0 site. The crystallographic screening did not pick up any of these binders. W08, however, binds in an unexpected orientation without occupying the Cap0 site or methionyl cavity. In all reported structures with ligands bound to the SAM site, the Cap0 binding site is occupied by one or two molecules of ethylene glycol used as cryogenic protectant for the crystals. It cannot be excluded that the short exposure of the cryoprotectant after the long compound soaking displaced potentially bound compounds although it is unlikely. Efforts have been made previously to find bi-substrate binders for nsp16. It does not seem trivial to bridge the gap between the SAM and Cap0 sites with linkers as is possible for nsp14 (Ahmed-Belkacem et al. 2020; Jung et al. 2022; Kottur et al. 2023). Since the extended pocket is adjacent to gate loop 2, compounds with large *N*7 extensions that populate this pocket and block the movement of gate loop 2 might be an alternative to bi-substrate inhibitors. This might prevent binding of the Cap0 substrate and, thereby, inhibit the enzyme function.

The mechanism of the Cap0 substrate binding and the simultaneous gate loop motions are not fully understood. For example, it has been unclear whether the occupation of the entire SAM site is needed to allow the gate loop movements necessary for binding Cap0-RNA which is observed in crystal structures including SAM and Cap0-RNA or the Cap0-analog (Minasov et al. 2021; Viswanathan et al. 2020; Wilamowski et al. 2021). Both loops shift by several Å upon binding of a Cap0 molecule. These dynamic changes are also observed in the present study when protein crystals were soaked with Cap0-analog or Cap0-RNA and SAM was replaced by sangivamycin or toyocamycin, which lack a methionyl moiety. The gate loop movements are very similar to what is observed in the SAM and Cap0-analog or Cap0-RNA, suggesting that ligand occupation of the methionyl cavity may not be a key factor for the gate loop motions.

Nsp10-16 contains multiple ligand binding sites but their dynamics, functions and potential for drug design are poorly understood. The phosphates of AMP, ADP, and ATP extend out of the SAM site near the position of the methionyl cavity. Due to the high flexibility of the phosphates, potential ligand-enzyme interactions are not clear. Nevertheless, this site could be explored for inhibitor design. Furthermore, as described recently, an additional cryptic pocket is located close to the extended *N*7 binding pocket. It could be exploited as a druggable site, e.g. via fragment merging, since covalent and non-covalent binders prevent occupation by SAM and inhibit enzyme activity (Inniss et al. 2023). We observed an EDTA molecule bound to the protein surface close to that position (figure S 6) which is also close to the *N*7-extension of W08. Another potential allosteric site with bound adenosine (Viswanathan et al. 2020) or ^m7^GTP (PDB entries 7JIB, 7JHE, 6WVN, 6WRZ) previously found in crystallographic studies has been suggested for drug targeting. This site is occupied by a buffer molecule (2-(*N*-morpholino)ethanesulfonic acid (MES)) in almost all structures reported here. These observations show that the potential ligand binding sites of nsp16 are still not fully understood and future studies will show their potential for drug discovery.

A significant complication concerning inhibitor discovery for nsp10-16 is that it is a multisubstrate enzyme. The binding-inhibition correlation is more complex than for single substrate enzymes and highly depends on the underlying kinetic mechanism (Sprenger et al. 2016; Yung-Chi and Prusoff 1973). Furthermore, the SAM binding site is more than 50 % occupied with SAM when purified from *E. coli* and the protein becomes unstable upon removal of bound SAM, which challenges binding studies. Our results show that adenosine derivatives such as the adenosine phosphates can partially replace SAM and that the extended pocket can be occupied by several *N*7 modified tubercidin derivatives. Such compounds should be explored further to gain higher affinity and selectivity of SAM-derived nsp16 inhibitors. Crystallographic screening is a very powerful tool to find new inhibitors and pharmacophore features especially for challenging targets such as nsp16. Further understanding the dynamics, regulation and kinetic mechanism of the nsp10-16 methyltransferase complex will provide crucial knowledge about the RNA processing of coronaviruses. Exploring the ligand binding sites presented and investigated herein will support drug development targeting SARS-CoV-2 and related viruses.

## Materials and Methods

### Protein expression, purification, and crystallization

Sequences of nsp10 and nsp16 (from polyprotein rep, 1a-1b uniprot ID: P0DTD1) were codon optimized for expression in *Escherichia coli* (*E. coli*) and synthesized and cloned into the pET-DUET-1 vector with a C-terminal Tobacco Etch Virus (TEV) protease cleavage site (ENLYFQG) and hexahistidine tag (His_6_) (GenScript). The expression construct was transfected into *E. coli* BL21 (DE3) pLyS (Promega, Madison, WI) and grown in Luria-Bertani broth (LB) supplemented with 50 µg/mL ampicillin and 130 µg/mL kanamycin overnight at 37 °C and 220 rpm. For inoculation of 1 L LB supplemented with ampicillin, 3 mL of the overnight culture were used. At OD_600_ = 1.8 - 2 expression was induced by addition of 0.4 mM isopropyl-β-D-thiogalactopyranosid (IPTG) and further incubated at 18 °C for 16 h at 220 rpm. Cells were harvested by centrifugation for 1 h at 5.000 rcf and frozen until purification or used directly for protein purification. Cell pellets were resuspended in lysis buffer (50 mM Tris HCl, 500 mM NaCl, 2 mM MgCl_2_, 25 mM imidazole, 1 mM Tris-(2-carboxyethyl)phosphine (TCEP), 5 % (v/v) glycerol, pH 8.3) supplemented with EDTA-free protease inhibitor mix, lysozyme, and DNase. Cell lysate was sonicated with 40 % amplitude for 5 s on, 25 s off cycle for 25 min at 4 °C and cleared by ultracentrifugation at 120 000 rcf at 4 °C for 45 min. The supernatant was collected and loaded onto a His-Trap FF column containing a Ni-NTA resin using a GE Healthcare ÄKTA Pure system with loading buffer (50 mM Tris HCl, 500 mM NaCl, 2 mM MgCl_2_, 25 mM, 1 mM TCEP, 5 % (v/v) glycerol, pH 8.3). The column was washed with loading buffer and protein eluted with elution buffer (50 mM Tris-HCl, 500 mM NaCl, 500 mM imidazole, pH 8.3). The pooled protein fractions were dialyzed overnight in dialysis buffer (50 mM Tris HCl, 500 mM NaCl, 2 mM MgCl_2_, 1 mM TCEP, 5 % (v/v) glycerol, pH 8.3) at 4 °C and His_6_-tag was removed by adding TEV protease (3 mg per pellet out of 3 L expression culture). The cleaved tag was separated from the protein by Ni-NTA affinity chromatography and the protein further purified by size exclusion chromatography (Superdex 200 16/600 column) in crystallization buffer (10 mM Tris HCs, 150 mM NaCl, 2 m MgCl_2_, 1 mM TCEP, 5 % (v/v) glycerol, pH 8.3). The protein was concentrated to 6 - 7 mg/mL (122 - 143 µM) and used immediately for crystallization or flash frozen in liquid nitrogen and stored at −80 °C.

### Crystallization and compound soaking

Protein crystals were grown by the hanging drop vapor diffusion method. For that, 1 µL protein with a concentration of 6.3 mg/mL and 1 µL reservoir solution (400 mM NaF, 100 mM MES at pH 6.5) were mixed. Rice shaped crystals of 75 – 100 µm length grew for approximately 19 days before they were soaked in reservoir solution supplemented with respective compounds. 5 – 20 crystals per compound were flash frozen in liquid nitrogen with 33 % ethylene glycol as cryogenic protectant. Compounds were dissolved to 100 mM in DMSO prior soaking and directly added to the drop in volumes that result in 10 mM ligand concentration in the crystal drop. For soaking with only one compound, the crystals were harvested 24 h after soaking. To obtain the structures of toyocamycin and sangivamycin in the presence of Cap-0-analog (^m7^GpppA purchased from New England Bioscience) or Cap0-RNA (Sequence: ^m7^GpppA UUAAA GGUUU AUACC UUCCC AGGUA, synthesized by BOCSCI INC, purified by HPLC, purity > 90 %), the compounds were first soaked for 24 h with 10 mM of tubercidin derivate and subsequently with 10 and 0.3 mM of Cap0 and Cap0-RNA, respectively.

### Selection of compounds for crystallographic screening

The entire list of compounds is documented in the supplementary material (table S 2). The selected compounds include a small compound library of SAM analogs purchased from ASINEX (80 compounds), known methyltransferase inhibitors, theophylline, tubercidin and adenine or adenosine derivatives as well as selected compounds from a docking study (figure S 4)

### Molecular docking and selection of adenosine derivatives from ASINEX purine library

The purchasable ASINEX compounds from the SAM analog, all nucleoside mimetics, and purine nucleoside libraries (3627 compounds) were prepared by UNICON (Sommer et al. 2016) using the best-scored protonation and tautomeric state per compound. We analyzed the crossdocking performance for the known SAM-site binders SAH, SAM, sinefungin and MTA, and the non-binders NTD008 and AZADC with an in-house version of the molecular docking tool JAMDA (Flachsenberg et al. 2023). To this end, we used the PDB entries 6W4H with SAM, 6W75 with SAM, 6WKQ with sinefungin, 6WQ3 with SAH, 6WRZ with SAH, 6WVN with SAM, 6XKM with SAM, 7JPE with SAM, 6WKS with SAM, 6YZ1 with sinefungin, 7BQ7 with SAM, 7C2I with SAM, 7C2J with SAM, and 7JYY with SAM. All structures were aligned to PDB entry 6W4H and the bound SAM was used as a reference ligand to define the binding site in a 6.5 Å radius. For the PDB entry 6YZ1, we observed the best cross-docking and ranking performance when retaining all water molecules considered as relevant in the docking preparation step with JAMDA. The molecular docking was performed with JAMDA default settings and 1000 conformations per compound. All compounds of the ASINEX library were docked in this manner and analyzed with PyMOL (Schrödinger and DeLano 2020). The best-scored 50 compounds were selected for crystal soaking experiments. In addition, 43 compounds with lower ranks were also selected based on visual inspection and a score at least similar to the one obtained for SAM. All selected compounds are considerably larger than known binders and extend into the pocket usually occupied by Cap0. They were purchased and tested for binding using soaking experiments.

### Diverse compound selection and selection based on similarity to SAM

In addition to the docking-based selection, we also selected a diverse set of 50 compounds based on the extended connectivity fingerprint (ECFP) with a diameter of 4 (ECFP4) (Rogers and Hahn 2010) and a distance-based k-Medoids clustering. The compounds assigned as medoids were purchased and tested for binding in soaking experiments. Additionally, the eight most similar compounds to SAM in terms of the ECFP4-based Tanimoto coefficient from the ASINEX set were selected for soaking experiments. All 2D similarity analyses were performed with KNIME [REF: 10.1145/1656274.1656280].

### Data collection, structure solution, and refinement

Data sets were collected at the PETRA III storage ring at Deutsches Elektronen-Synchrotron (DESY) in Hamburg at the P11 beamline and EMBL Hamburg beamlines P13 and P14. Data were processed using XDS (Kabsch 2010). As high-resolution limit of the data set, a CC_1/2_ above 0.3 – 0.5 was selected. The in-house developed data base AMARCORD (the AMARCORD system will be described in a future publication) was used for data tracking and organization as well as a framework to run further scripts for processing and structure refinement. Within AMARCORD, data were processed using DIMPLE (Wojdyr et al. 2013) and refined with phenix.refine (Afonine et al. 2012) using the structure model of PDB entry 7JIB. The pipeline used for this was in principle the same as used in the DESY SARS-CoV-2 screening campaign on MPro (Günther et al. 2021), just omitting the clustering and PanDDA-step. Hit identification was done by visually inspecting the 2 mFo − DFc and mFo − DFc difference density maps. For the final structures, the refinement was done using phenix.refine with a template structure based on PDB entry 7JIB and subsequent model building and refinement was done by alternating rounds with COOT (Emsley et al. 2010) and phenix.refine. Data collection and refinement statistics are provided in table S 1.

### Structural SAM site comparison

We generated a GeoMine (Diedrich et al. 2021) database of potential human methyltransferases using all entries of the PDB (accessed: January 24^th^, 2024) in a complex with SAM or SAH. Additionally, we used all PDB entries with the GO annotation 0032259 (biological process: methylation). These entries were restricted to protein from *Homo sapiens*. We applied SiteMine (Reim et al. 2024) to compare the SAM binding site of nsp10-16 (PDB entry 7R1U, ligand three-letter-code of the binding site-defining binder: 4IK) to DoGSite3 (Graef, Ehrt, and Rarey 2023)- predicted and ligand-based binding sites in the generated GeoMine database. The matches were ranked using the formula 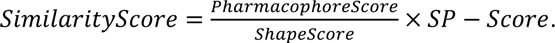 Default settings were used but the score normalization based on the larger pocket in terms of protein heavy atoms was disabled. The top-scored matches were visually inspected.

### Isothermal titration calorimetry

Affinities of compounds (Adenine, adenosine, AMP, ADP, ATP, MTA, 5-idotubercidin, toyocamycin, tubercidin, sangivamycin, sinefungin) to nsp10-16 were measured by ITC using a MicroCal PEAQ-ITC instrument (Malvern Panalytical). All titrations were performed at 25 °C, reference power of 10 µcal/s, stirring speed of 750 rpm, and a protein concentration of 45 µM in the sample cell and compound concentration of 500 µM in the syringe. All solutions contained the same buffer (50 mM Tris-HCl, pH 7.5, 200 mM NaCl). For each titration, one initial injection of 0.4 µL (0.8 s injection time) was followed by 12 or 18 injections with 2 µL each (4 s injection duration) and 150 s spacing between injections. Titrations of compound into buffer were performed as reference runs and subtracted from main runs. All thermograms were baseline-corrected and integrated using NITPIC (Keller et al. 2012). Data was fitted with a 1:1 binding model using SEDPHAT (H. Zhao, Piszczek, and Schuck 2015) and final plots were generated with GUSSI (Brautigam et al. 2016).

For the ITC displacement assay, 2 mM Sinefungin was titrated into 42.5 µM nsp10-16 mixed in advance with 2 mM sangivamycin. The buffer included 1 % DMSO. The K_D_ value for binding of sangivamycin was calculated with *ITCcalc* (Hammerschmidt et al. 2024).

### In vitro antiviral activity and cytotoxicity assays

Adenine, adenosine, AMP, ADP, ATP, MTA, sinefungin, toyocamycin, 5-idotubercidin, tubercidin, sangivamycin, and remdesivir were dissolved to 50 or 25 mM stock concentrations in either 100 % DMSO or water and stored at −20 °C. Vero E6 cells (ATCC CRL-1586) were grown in DMEM (PAN P04-03550) supplemented with 5 % FBS (PAN P30-3306), 1 mM sodium pyruvate (PAN P04-43100) and 1x antibiotic-antimycotic mix (ThermoFisher 15240062) at 37 °C and 5 % CO_2_.

Cells were seeded at 2 × 10^4^ cells/well in 96-well plates. After 24 h, the cell culture media was changed and serial dilutions of the compounds were added. Cell viability under 42 h compound treatment was determined via the Cell Counting Kit-8 (CCK-8, Sigma-Aldrich #96992) following the manufacturer’s instructions.

Cells seeded at 3.5 × 10^4^ cells/well in 96-well plates were pretreated 24 h later with serial dilutions of the compounds. After 1 h incubation with the compounds, SARS-CoV-2 (strain SARS-CoV-2/human/DEU/HH-1/2020) was subsequently added at a multiplicity of infection (MOI) of 0.01 and allowed absorption for 1 h. The viral inoculum was removed, cells were washed with PBS without Mg^2+^/Ca^2+^, and fresh media containing the compounds (final DMSO concentration 0.5 % (v/v)) was added to the cells. Cell culture supernatant was harvested 42 h post-infection and stored at −80 °C. Titers of infectious virus particles were measured via immunofocus assay. Briefly, cells seeded at 3.5 × 10^4^ cells/well in 96-well plates were inoculated with 50 µL of serial ten-fold dilutions of cell culture supernatant from treated cells. The inoculum was removed after 1 h and replaced by 1.5 % methylcellulose-DMEM (5 % FBS) overlay. Following incubation for 24 h, cells were inactivated and fixed with 4.5 % formaldehyde. Infected cells were detected using an antibody against SARS-CoV-2 nucleoprotein (ThermoFischer, PA5-81794). Foci were counted using an AID ELISpot reader from Mabtech. All experiments handling SARS-CoV-2 were performed at the biosafety level 3 (BSL-3) facilities in the Bernhard-Nocht Institute for Tropical Medicine in Hamburg under institutional safety guidelines.

The CC_50_ and EC_50_ values of the compounds were estimated by fitting the data to the sigmoidal four parameter logistic function with 1/Y^2^ weighting using GraphPad Prism version 9.5.1 for Mac OS X (GraphPad Software, San Diego, California USA, www.graphpad.com). The maximum values were fixed to 100 %, and the minimum values were constrained to be equal to or greater than the methodological detection limit. Samples deemed to be technical failures or extreme outliers were excluded from the calculations. The SI is defined as the ratio of CC_50_ over EC_50_.

### Microscale thermophoresis to investigate nsp10-16 ligand interactions

MST experiments were performed as described previously for FTAD displacement (Zimmermann et al. 2022) and Tris-NTA Red labeling (Schwickert et al. 2022). Briefly, His_6_-tagged nsp10-16 (expressed from HEK cells; commercially obtained from BPS Bioscience Catalog #100747) was labeled using the Monolith His-Tag Labeling Kit RED-Tris-NTA second generation (NanoTemper Technologies), according to the manufacturer’s instructions. Labeled protein was diluted to 10 nM into MST buffer (50 mM HEPES, 150 mM NaCl, 10 mM MgCl_2_, 0.05 % polysorbate-20, 0.1 % PEG-8000, pH 7.5), and the ligands were added in a final concentration of 1 mM from DMSO. Measurements were performed in triplets at 25 °C and at medium MST power in Monolith NT.115 Capillaries (NanoTemper Technologies). The raw data were analyzed using the MO.Affinity Analysis software (NanoTemper Technologies).

### NanoDSF ligand screening

Thermal shift assays were carried out in triplicate on the Prometheus NT.48 nanoDSF instrument (NanoTemper Technologies) using the manufacturer-designated capillaries. Sample solutions contained 5 µM protein and 10 mM of the investigated ligand in buffer (50 mM HEPES, 150 mM NaCl, 10 mM MgCl_2_, 0.05 % polysorbate-20, 0.1 % PEG-8000, pH 7.5) and 10 % (v/v) DMSO. In the capillaries, the sample solutions were heated from 20 °C to 80 °C at a heating rate of 1.5 °C/min, and fluorescence was recorded at 330 nm and 350 nm. The measured fluorescence ratio of the detected fluorescence was plotted as a function of temperature using GraphPad Prism 7.04. The melting temperature was calculated as the inflection point of the resulting sigmoidal curve.

### Spectral shift assay

Spectral shift assay was performed on a Monolith X instrument (NanoTemper Technologies) in parallel with MST measurements. Nsp10-16 purified from *E. coli* was labeled using the Monolith Lys-Tag Labeling Kit RED-NHS second generation (NanoTemper Technologies), according to the manufacturer’s instructions. Labeled protein was diluted to 25 nM into buffer (50 mM Tris-HCl, 200 mM NaCl, pH 8.3), and the ligands were added in a final concentration of 150 µM SAM and 250 µM Cap0-analog. Measurements were performed at 25 °C and at medium MST power in Monolith Premium Capillaries (NanoTemper Technologies). The raw data were analyzed using the MO.Affinity Analysis software (NanoTemper Technologies).

## Acknowledgments

We acknowledge Deutsches Elektronen-Synchrotron (DESY), Hamburg, Germany, a member of the Helmholtz Association HGF, for the provision of experimental facilities. Parts of this research were carried out at PETRA III and we would like to thank Johanna Hakanpää, Sofiane Saouane, Guillaume Pompidor, and Helena Taberman for assistance in using photon beamline P11. Beamtime was allocated for proposal I-20211542 and STP-20010439. Furthermore, synchrotron data was collected at beamlines P13 and P14 (proposal mx807) operated by EMBL Hamburg at the PETRA III storage ring (DESY, Hamburg, Germany). We would like to thank Gleb Bourenkov and Selina Storm for the assistance in using the beamline. We thank Dr. Spyros Chatziefthymiou for discussions on the experiments. This research was supported in part through the Maxwell computational resources operated at DESY.

## List of PDB entries

8BZV (ADN), 8OTO (AMP), 8OT0 (MTA+Gly), 8OV1 (ADP), 8OV2 (SGV), 8OV3 (5ID), 8OV4 (TO1), 8C5M (MTA), 8OTR (W08), 8OSX (ATP), 8BSD (TBN), 9EMV (SGV+Cap0), 8S8W (SGV+RNA), 8S8X (TO1+Cap0), 9EMJ (TO1+RNA), 9EML (Cap0+EDTA)

## References

Afonine, Pavel V, Ralf W Grosse-Kunstleve, Nathaniel Echols, Jeffrey J Headd, Nigel W Moriarty, Marat Mustyakimov, Thomas C Terwilliger, Alexandre Urzhumtsev, Peter H Zwart, and Paul D Adams. 2012. “Towards Automated Crystallographic Structure Refinement with Phenix. Refine.” Acta Crystallographica Section D: Biological Crystallography 68 (4): 352–67.

Ahmed-Belkacem, Rostom, Priscila Sutto-Ortiz, Mathis Guiraud, Bruno Canard, Jean-Jacques Vasseur, Etienne Decroly, and Françoise Debart. 2020. “Synthesis of Adenine Dinucleosides SAM Analogs as Specific Inhibitors of SARS-CoV Nsp14 RNA Cap Guanine-N7-Methyltransferase.” European Journal of Medicinal Chemistry 201: 112557.

Berg, Hannes, Maria A Wirtz Martin, Nadide Altincekic, Islam Alshamleh, Jasleen Kaur Bains, Julius Blechar, Betül Ceylan, et al. 2022. “Comprehensive Fragment Screening of the SARS-CoV-2 Proteome Explores Novel Chemical Space for Drug Development.” Angewandte Chemie 134 (46): e202205858.

Bergant, Valter, Shintaro Yamada, Vincent Grass, Yuta Tsukamoto, Teresa Lavacca, Karsten Krey, Maria-Teresa Mühlhofer, et al. 2022. “Attenuation of SARS-CoV-2 Replication and Associated Inflammation by Concomitant Targeting of Viral and Host Cap 2’-O-Ribose Methyltransferases.” The EMBO Journal 41 (17): e111608.

Bergstrom, Donald E, Alan J Brattesani, Mark K Ogawa, P Anantha Reddy, Michael J Schweickert, Jan Balzarini, and Erik De Clercq. 1984. “Antiviral Activity of C-5 Substituted Tubercidin Analogs.” Journal of Medicinal Chemistry 27 (3): 285–92.

Bernal, A Jayk, MM Gomes da Silva, DB Musungaie, E Kovalchuk, A Gonzalez, V Delos Reyes, A Martín-Quirós, et al. 2022. “Molnupiravir for Oral Treatment of COVID-19 in Nonhospitalized Patients., 2022” New England Journal of Medicine 386 (6): 509–20.

Bobileva, Olga, Raitis Bobrovs, Iveta Kanepe, Liene Patetko, Gints Kalnins, Mihails Sisovs, Anna L Bula, et al. 2021. “Potent SARS-CoV-2 mRNA Cap Methyltransferase Inhibitors by Bioisosteric Replacement of Methionine in SAM Cosubstrate.” ACS Medicinal Chemistry Letters 12 (7): 1102–7.

Bouvet, Mickaël, Claire Debarnot, Isabelle Imbert, Barbara Selisko, Eric J Snijder, Bruno Canard, and Etienne Decroly. 2010. “In Vitro Reconstitution of SARS-Coronavirus mRNA Cap Methylation.” PLoS Pathog 6 (4): e1000863.

Bradrick, Shelton S. 2017. “Causes and Consequences of Flavivirus RNA Methylation.” Frontiers in Microbiology 8: 2374.

Brautigam, Chad A, Huaying Zhao, Carolyn Vargas, Sandro Keller, and Peter Schuck. 2016. “Integration and Global Analysis of Isothermal Titration Calorimetry Data for Studying Macromolecular Interactions.” Nature Protocols 11 (5): 882–94.

Byszewska, Magdalena, Miros\law Śmietański, Elżbieta Purta, and Janusz M Bujnicki. 2014. “RNA Methyltransferases Involved in 5’ Cap Biosynthesis.” RNA Biology 11 (12): 1597–1607.

Chang, Li-Jen, and Tsung-Hsien Chen. 2021. “NSP16 2′-O-MTase in Coronavirus Pathogenesis: Possible Prevention and Treatments Strategies.” Viruses 13 (4): 538.

Chatterjee, Srijan, Manojit Bhattacharya, Kuldeep Dhama, Sang-Soo Lee, and Chiranjib Chakraborty. 2023. “Resistance to Nirmatrelvir Due to Mutations in the Mpro in the Subvariants of SARS-CoV-2 Omicron: Another Concern?” Molecular Therapy-Nucleic Acids 32: 263–66.

Chen, Yu, Hui Cai, Nian Xiang, Po Tien, Tero Ahola, Deyin Guo, and others. 2009. “Functional Screen Reveals SARS Coronavirus Nonstructural Protein Nsp14 as a Novel Cap N7 Methyltransferase.” Proceedings of the National Academy of Sciences 106 (9): 3484–89.

Chen, Yu, and Deyin Guo. 2016. “Molecular Mechanisms of Coronavirus RNA Capping and Methylation.” Virologica Sinica 31 (1): 3–11.

Chen, Yu, Ceyang Su, Min Ke, Xu Jin, Lirong Xu, Zhou Zhang, Andong Wu, et al. 2011. “Biochemical and Structural Insights into the Mechanisms of SARS Coronavirus RNA Ribose 2’-O-Methylation by Nsp16/Nsp10 Protein Complex.” PLoS Pathogens 7 (10): e1002294.

Cougot, Nicolas, Erwin van Dijk, Sylvie Babajko, and Bertrand Séraphin. 2004. “Cap-Tabolism.” Trends in Biochemical Sciences 29 (8): 436–44.

Daffis, Stephane, Kristy J Szretter, Jill Schriewer, Jianqing Li, Soonjeon Youn, John Errett, Tsai-Yu Lin, et al. 2010. “2′-O Methylation of the Viral mRNA Cap Evades Host Restriction by IFIT Family Members.” Nature 468 (7322): 452–56.

De Clercq, Erik, R Bernaerts, DE Bergstrom, MJ Robins, JA Montgomery, and A Holy. 1986. “Antirhinovirus Activity of Purine Nucleoside Analogs.” Antimicrobial Agents and Chemotherapy 29 (3): 482–87.

Decroly, Etienne, Isabelle Imbert, Bruno Coutard, Mickaël Bouvet, Barbara Selisko, Karine Alvarez, Alexander E Gorbalenya, Eric J Snijder, and Bruno Canard. 2008. “Coronavirus Nonstructural Protein 16 Is a Cap-0 Binding Enzyme Possessing (Nucleoside-2’ O)-Methyltransferase Activity.” Journal of Virology 82 (16): 8071–84.

Devkota, Kanchan, Matthieu Schapira, Sumera Perveen, Aliakbar Khalili Yazdi, Fengling Li, Irene Chau, Pegah Ghiabi, et al. 2021. “Probing the SAM Binding Site of SARS-CoV-2 Nsp14 in Vitro Using SAM Competitive Inhibitors Guides Developing Selective Bisubstrate Inhibitors.” SLAS DISCOVERY: Advancing the Science of Drug Discovery 26 (9): 1200–1211.

Diedrich, Konrad, Joel Graef, Katrin Schöning-Stierand, and Matthias Rarey. 2021. “GeoMine: Interactive Pattern Mining of Protein–Ligand Interfaces in the Protein Data Bank.” Bioinformatics 37 (3): 424–25.

Diedrich, Konrad, Bennet Krause, Ole Berg, and Matthias Rarey. 2023. “PoseEdit: Enhanced Ligand Binding Mode Communication by Interactive 2D Diagrams.” Journal of Computer-Aided Molecular Design 37 (10): 491–503.

Dong, Hongping, Bo Zhang, and Pei-Yong Shi. 2008. “Flavivirus Methyltransferase: A Novel Antiviral Target.” Antiviral Research 80 (1): 1–10.

Dong, Yetian, Tong Dai, Yujun Wei, Long Zhang, Min Zheng, and Fangfang Zhou. 2020. “A Systematic Review of SARS-CoV-2 Vaccine Candidates.” Signal Transduction and Targeted Therapy 5 (1): 237.

El Hassab, Mahmoud A, Tamer M Ibrahim, Sara T Al-Rashood, Amal Alharbi, Razan O Eskandrani, and Wagdy M Eldehna. 2021. “In Silico Identification of Novel SARS-COV-2 2′-O-Methyltransferase (Nsp16) Inhibitors: Structure-Based Virtual Screening, Molecular Dynamics Simulation and MM-PBSA Approaches.” Journal of Enzyme Inhibition and Medicinal Chemistry 36 (1): 727–36.

Emsley, Paul, Bernhard Lohkamp, William G Scott, and Kevin Cowtan. 2010. “Features and Development of Coot.” Acta Crystallographica Section D: Biological Crystallography 66 (4): 486–501.

Eyer, Luděk, Markéta Šmídková, Radim Nencka, Jiří Neča, Tomáš Kastl, Martin Palus, Erik De Clercq, and Daniel Růžek. 2016. “Structure-Activity Relationships of Nucleoside Analogues for Inhibition of Tick-Borne Encephalitis Virus.” Antiviral Research 133: 119–29.

Ferreira de Freitas, Renato, Danton Ivanochko, and Matthieu Schapira. 2019. “Methyltransferase Inhibitors: Competing with, or Exploiting the Bound Cofactor.” Molecules 24 (24): 4492.

Flachsenberg, Florian, Christiane Ehrt, Torben Gutermuth, and Matthias Rarey. 2023. “Redocking the PDB.” Journal of Chemical Information and Modeling.

Food, and Drug Administration (FDA). 2022. “Drug Administration Coronavirus (COVID-19) Update: FDA Authorizes First Oral Antiviral for Treatment of COVID-19.” Press Release.

Food, and Druga Administration (FDA). 2021. “Emergency Use Authorization 105. Paxlovid (Nirmatrelvir Co-Packaged with Ritonavir) for the Treatment of Mild-to-Moderate Coronavirus Disease 2019 (COVID-19) in Certain Adults and Pediatric Patients. Dec 22, 2021.” Available from: https://www.fda.gov/media/155049/download.

Furuichi, Yasuhiro, and Aaron J Shatkin. 2000. “Viral and Cellular mRNA Capping: Past and Prospects.”

Garone, Caterina, Aaron R D’Souza, Cristina Dallabona, Tiziana Lodi, Pedro Rebelo-Guiomar, Joanna Rorbach, Maria Alice Donati, et al. 2017. “Defective Mitochondrial rRNA Methyltransferase MRM2 Causes MELAS-like Clinical Syndrome.” Human Molecular Genetics 26 (21): 4257–66.

Graef, Joel, Christiane Ehrt, and Matthias Rarey. 2023. “Binding Site Detection Remastered: Enabling Fast, Robust, and Reliable Binding Site Detection and Descriptor Calculation with DoGSite3.” Journal of Chemical Information and Modeling 63 (10): 3128–37.

Grant, Rebecca, Jilian A Sacks, Priya Abraham, Supamit Chunsuttiwat, Cheryl Cohen, J Peter Figueroa, Thomas Fleming, et al. 2023. “When to Update COVID-19 Vaccine Composition.” Nature Medicine 29 (4): 776–80.

Günther, Sebastian, Patrick YA Reinke, Yaiza Fernández-García, Julia Lieske, Thomas J Lane, Helen M Ginn, Faisal HM Koua, et al. 2021. “X-Ray Screening Identifies Active Site and Allosteric Inhibitors of SARS-CoV-2 Main Protease.” Science 372 (6542): 642–46.

Hammerschmidt, Stefan J, Fabian Barthels, Annabelle C Weldert, and Christian Kersten. 2024. “Advanced Isothermal Titration Calorimetry for Medicinal Chemists with ITCcalc.” Journal of Chemical Education.

Imprachim, Nergis, Yuliana Yosaatmadja, and Joseph A Newman. 2023. “Crystal Structures and Fragment Screening of SARS-CoV-2 NSP14 Reveal Details of Exoribonuclease Activation and mRNA Capping and Provide Starting Points for Antiviral Drug Development.” Nucleic Acids Research 51 (1): 475–87.

Inniss, Nicole L, Ján Kozic, Fengling Li, Monica Rosas-Lemus, George Minasov, Jiří Rybáček, Yingjie Zhu, et al. 2023. “Discovery of a Druggable, Cryptic Pocket in SARS-CoV-2 Nsp16 Using Allosteric Inhibitors.” ACS Infectious Diseases.

Jung, Eunkyung, Ruben Soto-Acosta, Jiashu Xie, Daniel J Wilson, Christine D Dreis, Ryuichi Majima, Tiffany C Edwards, Robert J Geraghty, and Liqiang Chen. 2022. “Bisubstrate Inhibitors of Severe Acute Respiratory Syndrome Coronavirus-2 Nsp14 Methyltransferase.” ACS Medicinal Chemistry Letters 13 (9): 1477–84.

Kabinger, Florian, Carina Stiller, Jana Schmitzová, Christian Dienemann, Goran Kokic, Hauke S Hillen, Claudia Höbartner, and Patrick Cramer. 2021. “Mechanism of Molnupiravir-Induced SARS-CoV-2 Mutagenesis.” Nature Structural & Molecular Biology 28 (9): 740–46.

Kabsch, Wolfgang. 2010. “XDS.” Acta Crystallographica Section D: Biological Crystallography 66 (2): 125–32.

Keller, Sandro, Carolyn Vargas, Huaying Zhao, Grzegorz Piszczek, Chad A Brautigam, and Peter Schuck. 2012. “High-Precision Isothermal Titration Calorimetry with Automated Peak-Shape Analysis.” Analytical Chemistry 84 (11): 5066–73.

Khalifa, Shaden AM, Nermeen Yosri, Mohamed F El-Mallah, Reem Ghonaim, Zhiming Guo, Syed Ghulam Musharraf, Ming Du, et al. 2021. “Screening for Natural and Derived Bio-Active Compounds in Preclinical and Clinical Studies: One of the Frontlines of Fighting the Coronaviruses Pandemic.” Phytomedicine 85: 153311.

Klima, Martin, Aliakbar Khalili Yazdi, Fengling Li, Irene Chau, Taraneh Hajian, Albina Bolotokova, H Ümit Kaniskan, et al. 2022. “Crystal Structure of SARS-CoV-2 Nsp10–Nsp16 in Complex with Small Molecule Inhibitors, SS148 and WZ16.” Protein Science 31 (9): e4395.

Kottur, Jithesh, Kris M White, M Luis Rodriguez, Olga Rechkoblit, Richard Quintana-Feliciano, Ahana Nayar, Adolfo García-Sastre, and Aneel K Aggarwal. 2023. “Structures of SARS-CoV-2 N7-Methyltransferase with DOT1L and PRMT7 Inhibitors Provide a Platform for New Antivirals.” PLoS Pathogens 19 (7): e1011546.

Kozielski, Frank, Céleste Sele, Vladimir O Talibov, Jiaqi Lou, Danni Dong, Qian Wang, Xinyue Shi, et al. 2022. “Identification of Fragments Binding to SARS-CoV-2 Nsp10 Reveals Ligand-Binding Sites in Conserved Interfaces between Nsp10 and Nsp14/Nsp16.” RSC Chemical Biology 3 (1): 44–55.

Krafcikova, Petra, Jan Silhan, Radim Nencka, and Evzen Boura. 2020. “Structural Analysis of the SARS-CoV-2 Methyltransferase Complex Involved in RNA Cap Creation Bound to Sinefungin.” Nature Communications 11 (1): 3717.

Kuzikov, Maria, Elisa Costanzi, Jeanette Reinshagen, Francesca Esposito, Laura Vangeel, Markus Wolf, Bernhard Ellinger, et al. 2021. “Identification of Inhibitors of SARS-CoV-2 3CL-pro Enzymatic Activity Using a Small Molecule in Vitro Repurposing Screen.” ACS Pharmacology & Translational Science 4 (3): 1096–1110.

Lewis, Joe D, and Elisa Izaurflde. 1997. “The Role of the Cap Structure in RNA Processing and Nuclear Export.” European Journal of Biochemistry 247 (2): 461–69.

Li, Fengling, Pegah Ghiabi, Taraneh Hajian, Martin Klima, Alice Shi Ming Li, Aliakbar Khalili Yazdi, Irene Chau, et al. 2023. “SS148 and WZ16 Inhibit the Activities of Nsp10-Nsp16 Complexes from All Seven Human Pathogenic Coronaviruses.” Biochimica et Biophysica Acta (BBA)-General Subjects, 130319.

Lin, Sheng, Hua Chen, Fei Ye, Zimin Chen, Fanli Yang, Yue Zheng, Yu Cao, Jingxin Qiao, Shengyong Yang, and Guangwen Lu. 2020. “Crystal Structure of SARS-CoV-2 Nsp10/Nsp16 2’-O-Methylase and Its Implication on Antiviral Drug Design.” Signal Transduction and Targeted Therapy 5 (1): 1–4.

Liu, Lihui, Hongping Dong, Hui Chen, Jing Zhang, Hua Ling, Zhong Li, Pei-Yong Shi, and Hongmin Li. 2010. “Flavivirus RNA Cap Methyltransferase: Structure, Function, and Inhibition.” Frontiers in Biology 5: 286–303.

Malone, Brandon, Nadya Urakova, Eric J Snijder, and Elizabeth A Campbell. 2022. “Structures and Functions of Coronavirus Replication–Transcription Complexes and Their Relevance for SARS-CoV-2 Drug Design.” Nature Reviews Molecular Cell Biology 23 (1): 21–39.

Manandhar, Anjela, Benjamin E Blass, Dennis J Colussi, Imane Almi, Magid Abou-Gharbia, Michael L Klein, and Khaled M Elokely. 2021. “Targeting SARS-CoV-2 M3CLpro by HCV NS3/4a Inhibitors: In Silico Modeling and in Vitro Screening.” Journal of Chemical Information and Modeling 61 (2): 1020–32.

Maurya, Santosh K, Akhilesh Kumar Maurya, Nidhi Mishra, and Hifzur R Siddique. 2020. “Virtual Screening, ADME/T, and Binding Free Energy Analysis of Anti-Viral, Anti-Protease, and Anti-Infectious Compounds against NSP10/NSP16 Methyltransferase and Main Protease of SARS CoV-2.” Journal of Receptors and Signal Transduction 40 (6): 605–12.

Maveyraud, Laurent, and Lionel Mourey. 2020. “Protein X-Ray Crystallography and Drug Discovery.” Molecules 25 (5): 1030.

Menachery, Vineet D, Kari Debbink, and Ralph S Baric. 2014. “Coronavirus Non-Structural Protein 16: Evasion, Attenuation, and Possible Treatments.” Virus Research 194: 191–99.

Minasov, George, Monica Rosas-Lemus, Ludmilla Shuvalova, Nicole L Inniss, Joseph S Brunzelle, Courtney M Daczkowski, Paul Hoover, Andrew D Mesecar, and Karla JF Satchell. 2021. “Mn2+ Coordinates Cap-0-RNA to Align Substrates for Efficient 2’-O-Methyl Transfer by SARS-CoV-2 Nsp16.” Science signaling, 14 (689), eabh2071.

Mudgal, Rajat, Supreeti Mahajan, and Shailly Tomar. 2020. “Inhibition of Chikungunya Virus by an Adenosine Analog Targeting the SAM-Dependent nsP1 Methyltransferase.” FEBS Letters 594 (4): 678–94.

Otava, Tomas, Michal Sala, Fengling Li, Jindrich Fanfrlik, Kanchan Devkota, Sumera Perveen, Irene Chau, et al. 2021. “The Structure-Based Design of SARS-CoV-2 Nsp14 Methyltransferase Ligands Yields Nanomolar Inhibitors.” ACS Infectious Diseases 7 (8): 2214–20.

Pachetti, Maria, Bruna Marini, Francesca Benedetti, Fabiola Giudici, Elisabetta Mauro, Paola Storici, Claudio Masciovecchio, et al. 2020. “Emerging SARS-CoV-2 Mutation Hot Spots Include a Novel RNA-Dependent-RNA Polymerase Variant.” Journal of Translational Medicine 18: 1–9.

Parums, Dinah V. 2022. “Current Status of Oral Antiviral Drug Treatments for SARS-CoV-2 Infection in Non-Hospitalized Patients.” Medical Science Monitor: International Medical Journal of Experimental and Clinical Research 28: e935952–1.

Parums, Dinah V. 2023. “Factors Driving New Variants of SARS-CoV-2, Immune Escape, and Resistance to Antiviral Treatments as the End of the COVID-19 Pandemic Is Declared.” Medical Science Monitor: International Medical Journal of Experimental and Clinical Research 29: e942960–1.

Perveen, Sumera, Aliakbar Khalili Yazdi, Kanchan Devkota, Fengling Li, Pegah Ghiabi, Taraneh Hajian, Peter Loppnau, Albina Bolotokova, and Masoud Vedadi. 2021. “A High-Throughput RNA Displacement Assay for Screening SARS-CoV-2 Nsp10-Nsp16 Complex toward Developing Therapeutics for COVID-19.” SLAS DISCOVERY: Advancing the Science of Drug Discovery 26 (5): 620–27.

Pletzer-Zelgert, Jonathan, Christiane Ehrt, Inken Fender, Axel Griewel, Florian Flachsenberg, Gerhard Klebe, and Matthias Rarey. 2023. “LifeSoaks: A Tool for Analyzing Solvent Channels in Protein Crystals and Obstacles for Soaking Experiments.” Acta Crystallographica Section D: Structural Biology 79 (9).

Ramanathan, Anand, G Brett Robb, and Siu-Hong Chan. 2016. “mRNA Capping: Biological Functions and Applications.” Nucleic Acids Research 44 (16): 7511–26.

Reim, Thorben, Christiane Ehrt, Joel Graef, Sebastian Günther, Alke Meents, and Matthias Rarey. 2024. “SiteMine: Large-Scale Binding Site Similarity Searching in Protein Structure Databases.” *Archiv Der Pharmazie*, e2300661.

Rogers, David, and Mathew Hahn. 2010. “Extended-Connectivity Fingerprints.” Journal of Chemical Information and Modeling 50 (5): 742–54.

Rosas-Lemus, Monica, George Minasov, Ludmilla Shuvalova, Nicole L Inniss, Olga Kiryukhina, Joseph Brunzelle, and Karla JF Satchell. 2020. “High-Resolution Structures of the SARS-CoV-2 2’-O-Methyltransferase Reveal Strategies for Structure-Based Inhibitor Design.” Science Signaling 13 (651).

Saha, Abinit, Ashish Ranjan Sharma, Manojit Bhattacharya, Garima Sharma, Sang-Soo Lee, and Chiranjib Chakraborty. 2020. “Probable Molecular Mechanism of Remdesivir for the Treatment of COVID-19: Need to Know More.” Archives of Medical Research 51 (6): 585–86.

Sandmann, Frank G, Nicholas G Davies, Anna Vassall, W John Edmunds, Mark Jit, Fiona Yueqian Sun, C Julian Villabona-Arenas, et al. 2021. “The Potential Health and Economic Value of SARS-CoV-2 Vaccination alongside Physical Distancing in the UK: A Transmission Model-Based Future Scenario Analysis and Economic Evaluation.” The Lancet Infectious Diseases 21 (7): 962–74.

Schöning-Stierand, Katrin, Konrad Diedrich, Christiane Ehrt, Florian Flachsenberg, Joel Graef, Jochen Sieg, Patrick Penner, Martin Poppinga, Annett Ungethüm, and Matthias Rarey. 2022. “Proteins Plus: A Comprehensive Collection of Web-Based Molecular Modeling Tools.” Nucleic Acids Research 50 (W1): W611–15.

Schrödinger, LLC, and Warren DeLano. 2020. “PyMOL.” http://www.pymol.org/pymol.

Schultz, David C, Robert M Johnson, Kasirajan Ayyanathan, Jesse Miller, Kanupriya Whig, Brinda Kamalia, Mark Dittmar, et al. 2022. “Pyrimidine Inhibitors Synergize with Nucleoside Analogues to Block SARS-CoV-2.” Nature 604 (7904): 134–40.

Schwer, Beate, Xiangdong Mao, and Stewart Shuman. 1998. “Accelerated mRNA Decay in Conditional Mutants of Yeast mRNA Capping Enzyme.” Nucleic Acids Research 26 (9): 2050–57.

Schwickert, Marvin, Tim R Fischer, Robert A Zimmermann, Sabrina N Hoba, J Laurenz Meidner, Marlies Weber, Moritz Weber, et al. 2022. “Discovery of Inhibitors of DNA Methyltransferase 2, an Epitranscriptomic Modulator and Potential Target for Cancer Treatment.” Journal of Medicinal Chemistry 65 (14): 9750–88.

Sommer, Kai, Nils-Ole Friedrich, Stefan Bietz, Matthias Hilbig, Therese Inhester, and Matthias Rarey. 2016. “UNICON: A Powerful and Easy-to-Use Compound Library Converter.” Journal of Chemical Information and Modeling 56 (6): 1105–1111.

Sprenger, Janina, Jannette Carey, Bo Svensson, Verena Wengel, and Lo Persson. 2016. “Binding and Inhibition of Spermidine Synthase from Plasmodium Falciparum and Implications for in Vitro Inhibitor Testing.” Plos One 11 (9): e0163442.

Vijayan, Viswanathan, Pradeep Pant, Naval Vikram, Punit Kaur, TP Singh, Sujata Sharma, and Pradeep Sharma. 2020. “Identification of Promising Drug Candidates against NSP16 of SARS-CoV-2 through Computational Drug Repurposing Study.” Journal of Biomolecular Structure and Dynamics, 1–15.

Viswanathan, Thiruselvam, Shailee Arya, Siu-Hong Chan, Shan Qi, Nan Dai, Anurag Misra, Jun-Gyu Park, et al. 2020. “Structural Basis of RNA Cap Modification by SARS-CoV-2.” Nature Communications 11 (1): 1–7.

Walker, Alexander P, Haitian Fan, Jeremy R Keown, Jonathan M Grimes, and Ervin Fodor. 2021. “Identification of Guanylyltransferase Activity in the SARS-CoV-2 RNA Polymerase.” bioRxiv, 2021–03.

Wang, Zhonglei, and Liyan Yang. 2020. “Turning the Tide: Natural Products and Natural-Product-Inspired Chemicals as Potential Counters to SARS-CoV-2 Infection.” Frontiers in Pharmacology 11: 1013.

Werner, Maria, Elżbieta Purta, Katarzyna H Kaminska, Iwona A Cymerman, David A Campbell, Bidyottam Mittra, Jesse R Zamudio, Nancy R Sturm, Jacek Jaworski, and Janusz M Bujnicki. 2011. “2′-O-Ribose Methylation of Cap2 in Human: Function and Evolution in a Horizontally Mobile Family.” Nucleic Acids Research 39 (11): 4756–68.

Wilamowski, Mateusz, Darren A Sherrell, George Minasov, Youngchang Kim, Ludmilla Shuvalova, Alex Lavens, Ryan Chard, et al. 2021. “2′-O Methylation of RNA Cap in SARS-CoV-2 Captured by Serial Crystallography.” Proceedings of the National Academy of Sciences 118 (21): e2100170118.

Wojdyr, Marcin, Ronan Keegan, Graeme Winter, and Alun Ashton. 2013. “DIMPLE-a Pipeline for the Rapid Generation of Difference Maps from Protein Crystals with Putatively Bound Ligands.” Acta Crystallographica Section A 69: s299–s299.

World Health Organization 2023 data.who.int, WHO Coronavirus (COVID-19) dashboard > Deaths [Dashboard]. https://data.who.int/dashboards/covid19/deaths, accessed on 2023-12-15

Xu, Chi, Zunhui Ke, Chuandong Liu, Zhihao Wang, Denghui Liu, Lei Zhang, Jingning Wang, et al. 2020. “Systemic in Silico Screening in Drug Discovery for Coronavirus Disease (COVID-19) with an Online Interactive Web Server.” Journal of Chemical Information and Modeling 60 (12): 5735–45.

Yan, Liming, Ji Ge, Litao Zheng, Ying Zhang, Yan Gao, Tao Wang, Yucen Huang, et al. 2021. “Cryo-EM Structure of an Extended SARS-CoV-2 Replication and Transcription Complex Reveals an Intermediate State in Cap Synthesis.” Cell 184 (1): 184–93.

Yazdi, Aliakbar Khalili, Fengling Li, Kanchan Devkota, Sumera Perveen, Pegah Ghiabi, Taraneh Hajian, Albina Bolotokova, and Masoud Vedadi. 2021. “A High-Throughput Radioactivity-Based Assay for Screening SARS-CoV-2 Nsp10-Nsp16 Complex.” Slas Discovery 26 (6): 757–65.

Yung-Chi, Cheng, and William H Prusoff. 1973. “Relationship between the Inhibition Constant (KI) and the Concentration of Inhibitor Which Causes 50 per Cent Inhibition (I50) of an Enzymatic Reaction.” Biochemical Pharmacology 22 (23): 3099–3108.

Zhao, Huaying, Grzegorz Piszczek, and Peter Schuck. 2015. “SEDPHAT–a Platform for Global ITC Analysis and Global Multi-Method Analysis of Molecular Interactions.” Methods 76: 137–48.

Zhao, Jianyuan, Qian Liu, Dongrong Yi, Quanjie Li, SaiSai Guo, Ling Ma, Yongxin Zhang, et al. 2022. “5-Iodotubercidin Inhibits SARS-CoV-2 RNA Synthesis.” Antiviral Research 198: 105254.

Zhou, Peng, Xing-Lou Yang, Xian-Guang Wang, Ben Hu, Lei Zhang, Wei Zhang, Hao-Rui Si, et al. 2020. “A Pneumonia Outbreak Associated with a New Coronavirus of Probable Bat Origin.” Nature 579 (7798): 270–73.

Zhu, Na, Dingyu Zhang, Wenling Wang, Xingwang Li, Bo Yang, Jingdong Song, Xiang Zhao, et al. 2020. “A Novel Coronavirus from Patients with Pneumonia in China, 2019.” New England Journal of Medicine 382: 727–733.

Zimmermann, Robert Alexander, Marvin Schwickert, J Laurenz Meidner, Zarina Nidoieva, Mark Helm, and Tanja Schirmeister. 2022. “An Optimized Microscale Thermophoresis Method for High-Throughput Screening of DNA Methyltransferase 2 Ligands.” ACS Pharmacology & Translational Science 5 (11): 1079–85.

