## Supplemental material for "SARS-CoV-2 methyltransferase nsp10-16 in complex with natural and drug-like purine analogs for guiding structure-based drug discovery"

### Supplementary

#### Supplementary results

##### FTAD binds to SAM site in crystal but not in solution

We tested the fluorescent probe 5-FAM-triazolyl-adenosyl-Dab (FTAD) together with in an effort to establish a microscale thermophoresis (MST) displacement assay for nsp10-16. Ligands that bind weakly to the SAM site would be displaced by SAM-like probes with an attached fluorophore which would result in a change of signal that can be translated into binding affinities (Zimmermann et al. 2022). The method seemed promising since FTAD was shown to bind to monomeric nsp16 and nsp14 (Samrat et al. 2023) in context of an MST displacement assay previously and to bind to several other methyltransferases even with $K_D$ values as high as 500 µM (Luan et al. 2016).

FTAD binds to nsp10-16 in the MST assay with an affinity of 1.17 µM in agreement with the report of Samrat et al. 2023 (figure S 6 A). However, attempts to displace the bound probe by a competitor such as SAM or SAH failed, suggesting that binding of FTAD leads to an irreversible change in protein conformation in the nsp10-16 complex, which is supported by the crystal structure we obtained in parallel. Ambiguous electron density was present in crystals soaked with FTAD which cannot be explained by SAM binding only (figure S 7). The SAM-like part seems to bind similarly to the substrate SAM. The fluorophore must be very flexible and extends in the direction of the methionine moiety into the solvent without interacting with the protein (figure S 7) which would be beneficial for an MST displacement assay if protein integrity would not be affected. Adding 5 mM FTAD to nsp10-16 crystals for soaking experiments led to disintegration of crystals within seconds. A concentration of 0.6 mM was tolerated by the crystals, however, nsp10 seems to have lost a zinc ion in one of the two zinc binding site which leads to disorder in this region and could influence nsp16 stability. These findings subsequently render FTAD unsuitable as a probe for the dimeric protein complex.

Therefore, we followed a second assay strategy and labeled the N-terminal His-tag using a non-covalent fluorescent Tris-NTA Red dye analogous to a protocol developed for the nsp10-14 methyltransferase (Kozielski et al. 2022). The globally labeled protein complex was used to determine the affinity of SAH and the Cap0 analog to 6.82 µM and 52.9 µM, respectively (figure S 6 B, C). However, the tubercidin derived ligands showed no effect on the thermophoresis of the protein complex at ligand concentrations up to 1 mM, reflecting their low affinity (figure S 6 D).

##### EDTA bound to nsp16 surface close to SAM site

An observation from the SAM and Cap0-analog containing structure determined here is the presence of an EDTA molecule on the nsp16 surface close to the SAM site at α-helix D nearby the C-terminus.

EDTA was added during the soaking to stop the methyl transfer reaction (Viswanathan et al. 2021). The occupancy is 0.75 and the density is ambiguous (figure S 6 A, B). The interaction with the protein surface is weak and might involve two water molecules, Ala6914, Gly6946, Thr6949, Tyr6950, Gly6953, and an ethylene glycol that was used as cryogenic protectant (figure S 6 C). A cryptic binding pocket that can bind a pyrimidin-2-ol inhibitor just below this EDTA binding site has been identified previously and changes protein conformation upon ligand binding (Inniss et al. 2023). Although the interaction of EDTA with nsp16 seems weak it shows that the capacity of nsp16 to bind diverse molecules to different sites is not yet fully understood.

#### Supplementary Figures

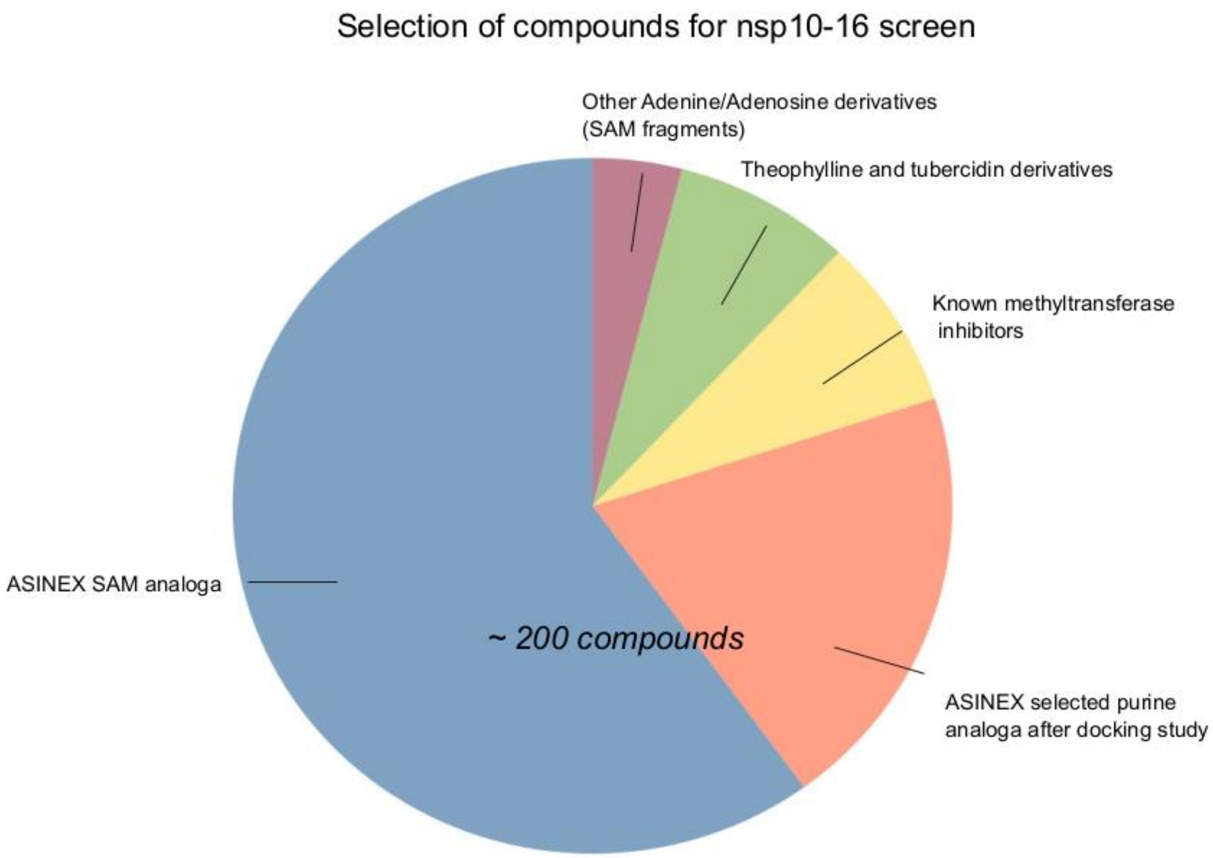


Figure S 1 ***Visual representation of compound selection for nsp10-16 binder X-ray screen.***

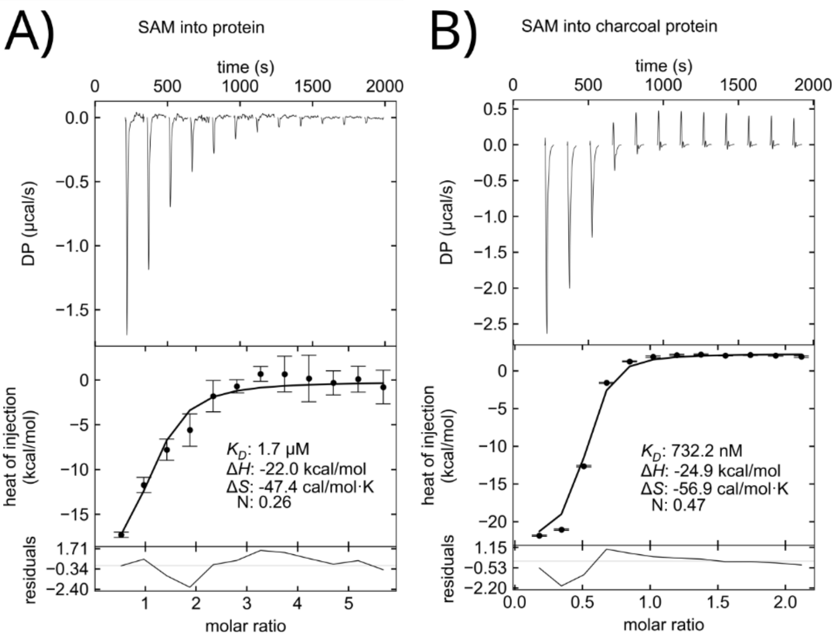


Figure S 2 **Isothermal titration calorimetry of purified nsp10-16.** A) The substrate SAM (500 µM) was titrated into nsp10-16 (45 µM) containing SAM from *E. coli* which was carried through the purification process. B) SAM was titrated into nsp10-16 that had been dialyzed extensively with charcoal.

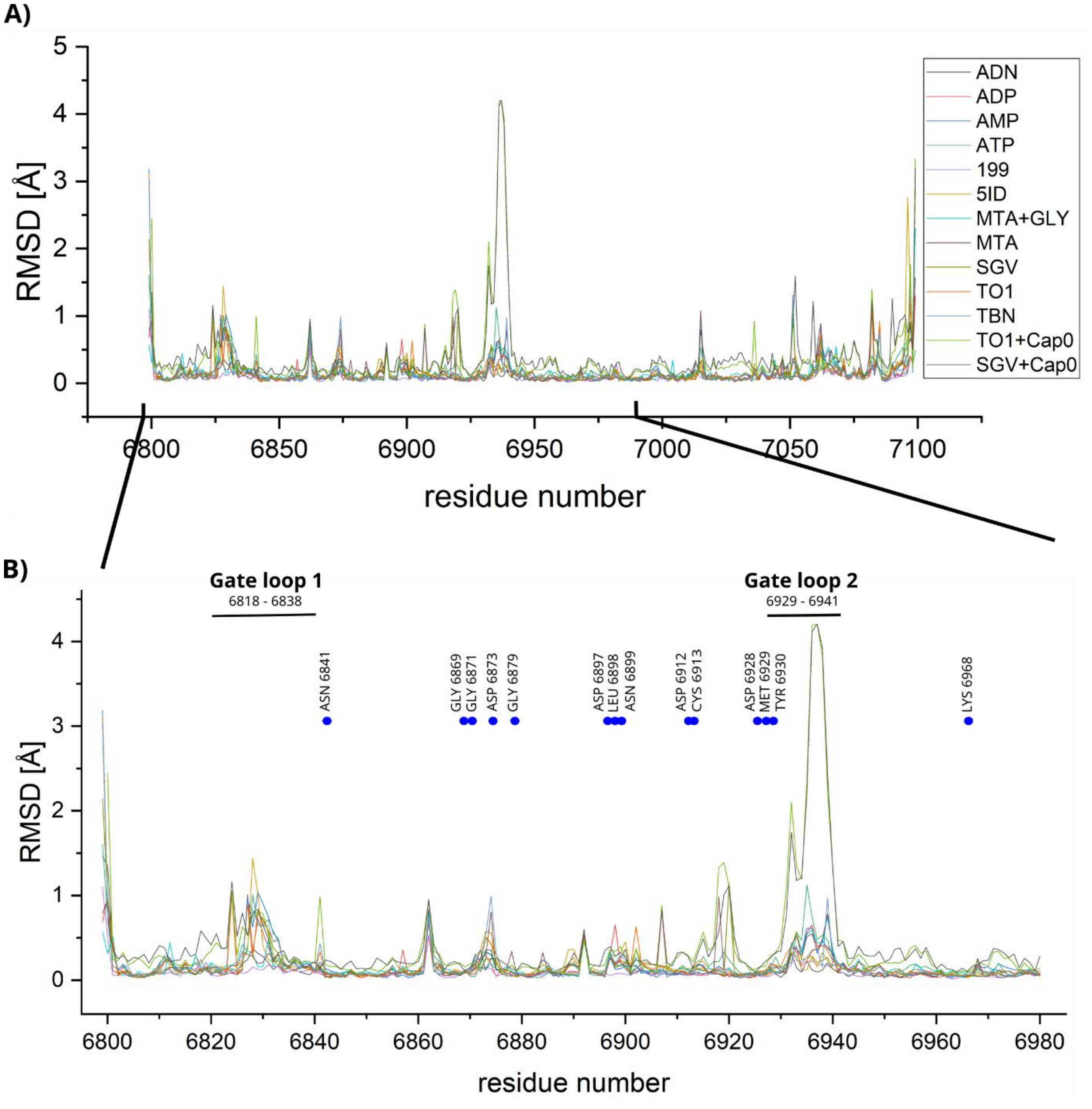


Figure S 3 **Backbone RMSD of all nsp16 residues with SAM bound structure as reference calculated with noc**. A) Overview of RMSD values for the complete nsp16 sequence. B) Nsp16 residues 6800 to 6980. Blue dots mark SAM interacting residues. Gate loop residues are indicated by a bar above.

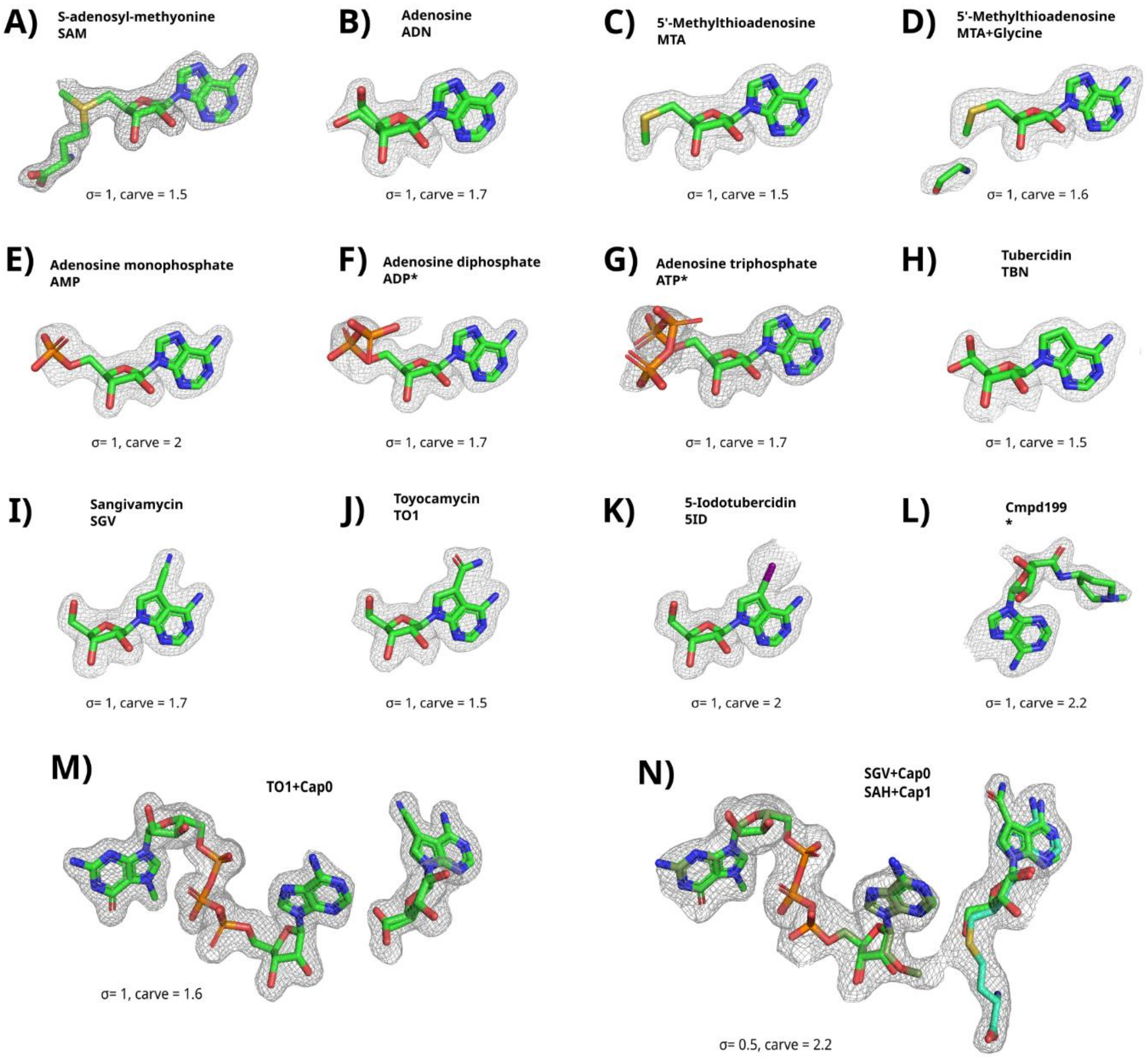


Figure S 4 **Compound electron densities in SAM binding site.**

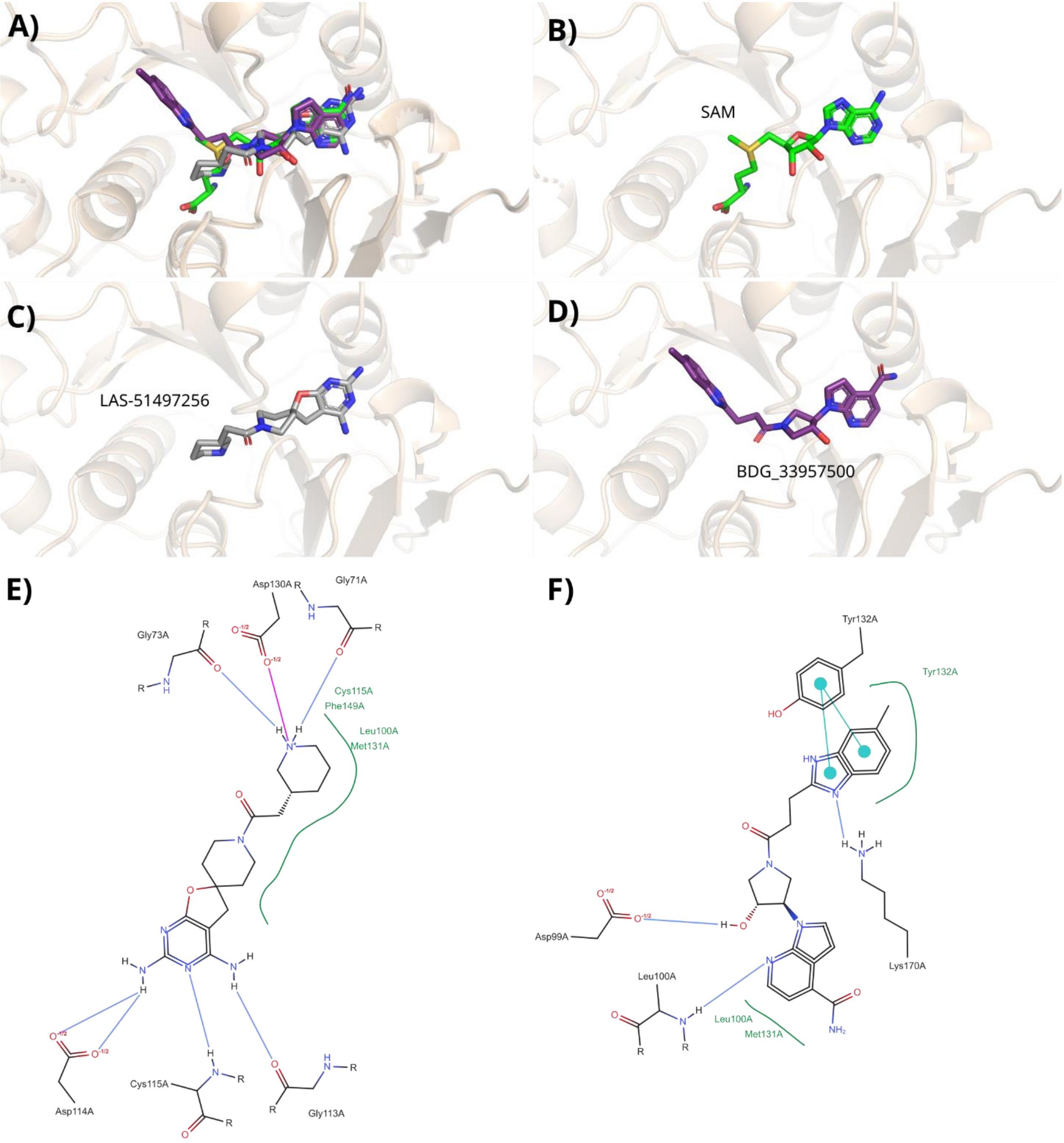


Figure S 5 **Docking results.** Examples of ligand poses for ligands covering only the SAM site or the SAM and the Cap0 site. A) Overlay of SAM with the docked ligands. B) SAM binding mode. C) LAS_51497256 (rank 11) binding mode that covers the SAM site. D) BDG_33957500 (rank 12) binding mode that covers the SAM and partially the Cap0 site. E) and F) Ligand-protein interactions of LAS_51497256 and BDG_33957500, respectively. Plots were created and edited with PoseView and PoseEdit ( https://proteins.plus, Schöning-Stierand et al. 2022; Diedrich et al. 2023).

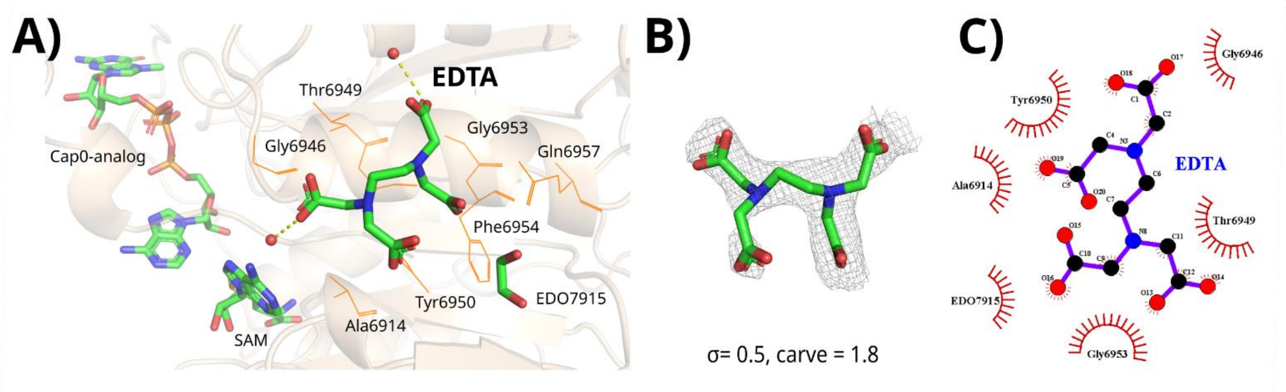


Figure S 6 **EDTA bound to nsp10-16 surface close to SAM binding site.** A) nsp16 are shown as orange cartoon; EDTA, SAM, and Cap0-analog are represented as green sticks; nsp16 residues that are close to EDTA are shown as lines. B) Electron density of EDTA (occupancy: 75 %). C) Interacting residues identified by LigPlot.

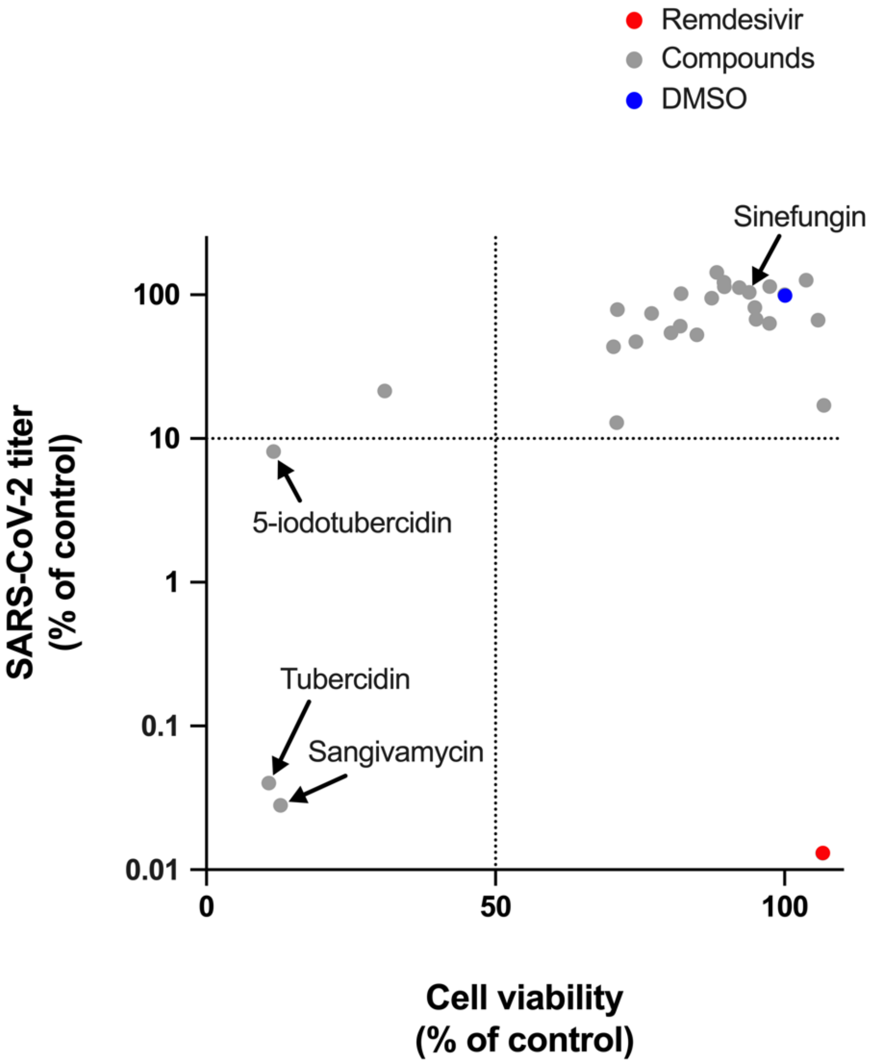


Figure S 7 **Biological relevance of SARS-CoV-2 nsp10/nsp16 complex crystallographic binders for in vitro virus replication.** Identified binders from crystallographic study (gray circles) and remdesivir (red circle), as a positive control of the virus replicative inhibition, were used at 10 μM concentrations to treat infected Vero E6 cells for 42 h. The solvent DMSO (blue circle) at 0.5% (v/v) was present in all conditions and used as the negative control. Viral titers and cell viability were determined via immunofocus assays and the CCK-8 method, respectively. The data displays the mean of three independent replicates in one experiment. The outcome of compounds, tubercidin, 5-iodotubercidin, sangivamycin, and sinefungin are labeled.

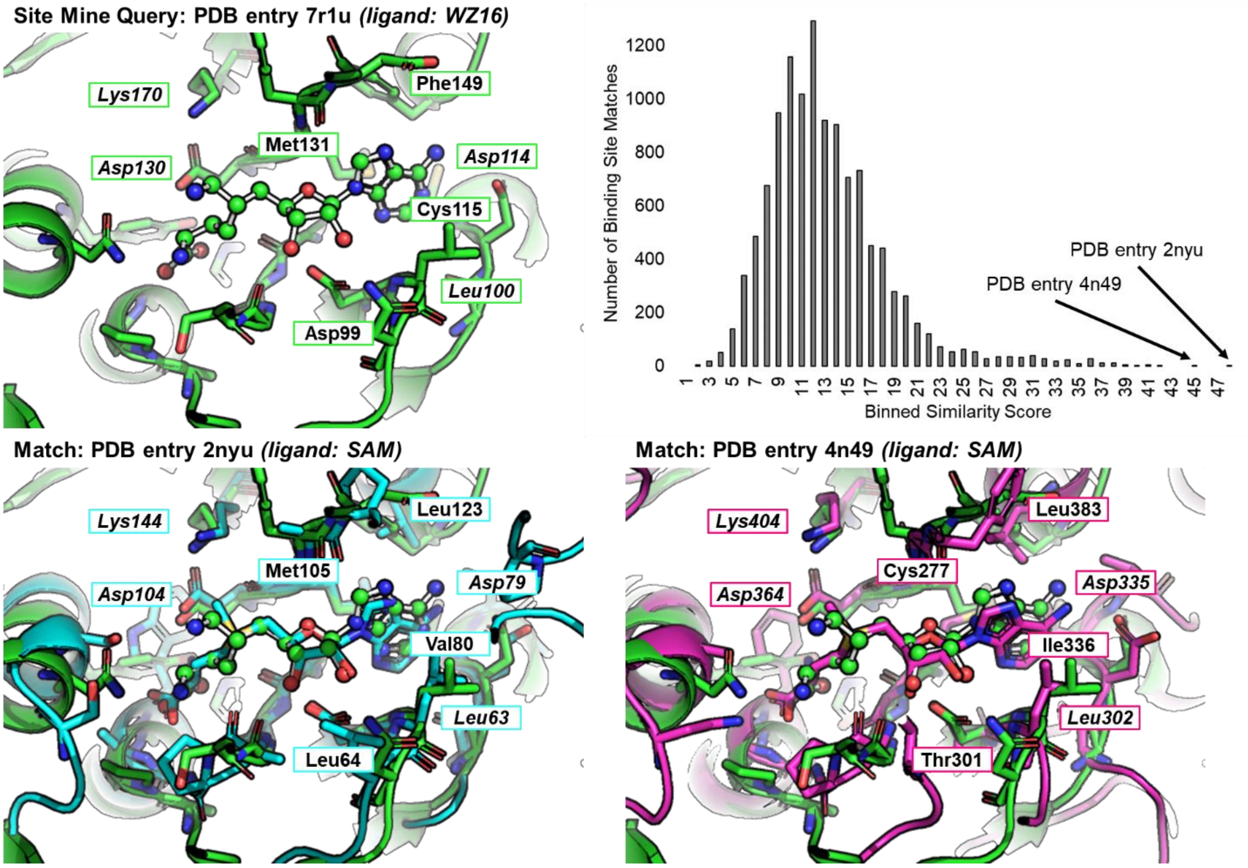


Figure S 8 **SiteMine results.** The most similar aligning residues in the SiteMine alignments are labeled by their residue name and ID. Residues in italics are conserved in all three binding sites.

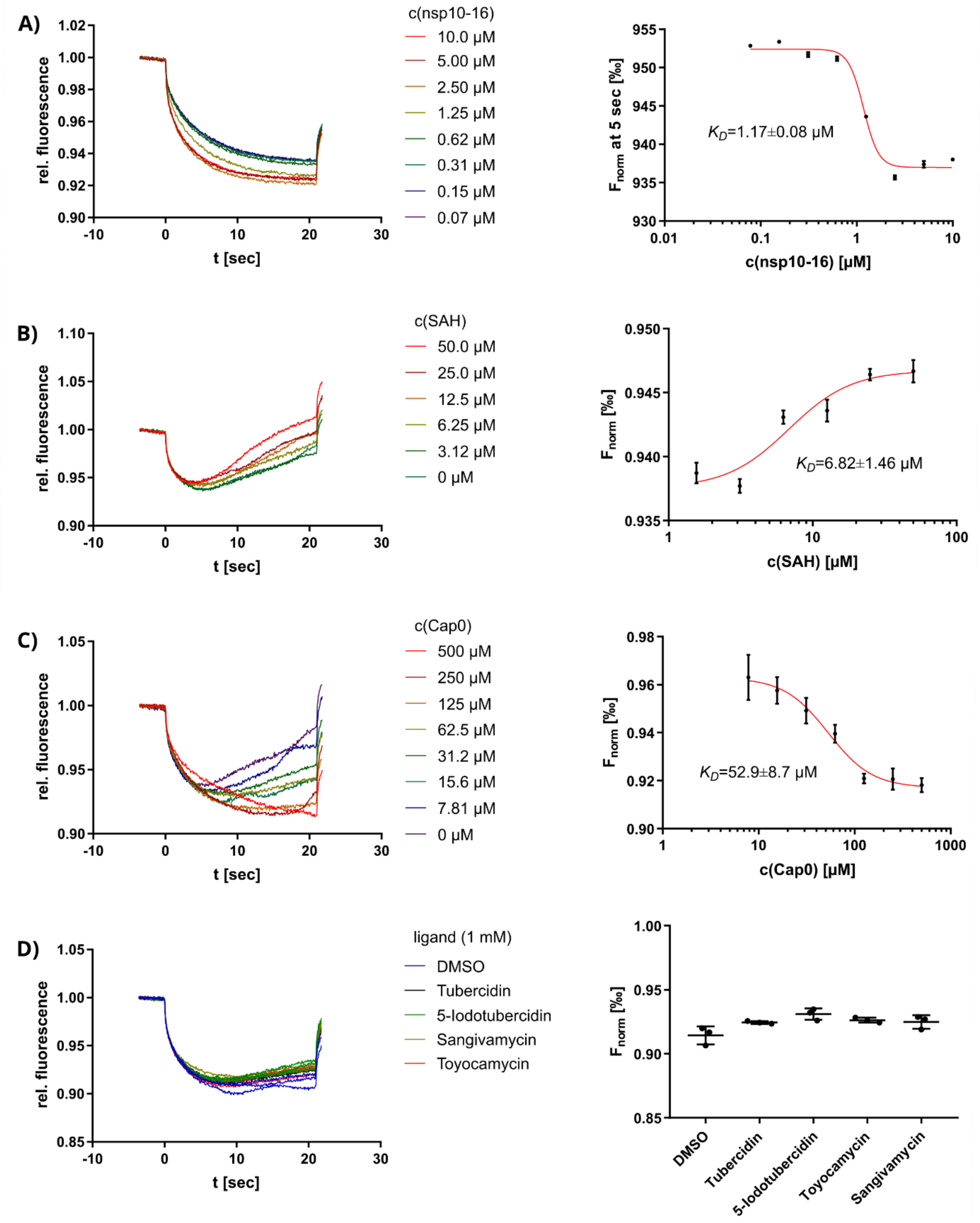


Figure S 9 **MST measurements for binders.** A) Affinity determination between FTAD and nsp10-16. (B) Affinity determination of SAH with Tris-NTA Red labeled nsp10-16. C) Affinity determination of the Cap0 analog with Tris-NTA Red labeled nsp10-16. D) Tubericidin-like ligand screening (1 mM) on Tris-NTA Red labeled nsp10-16.

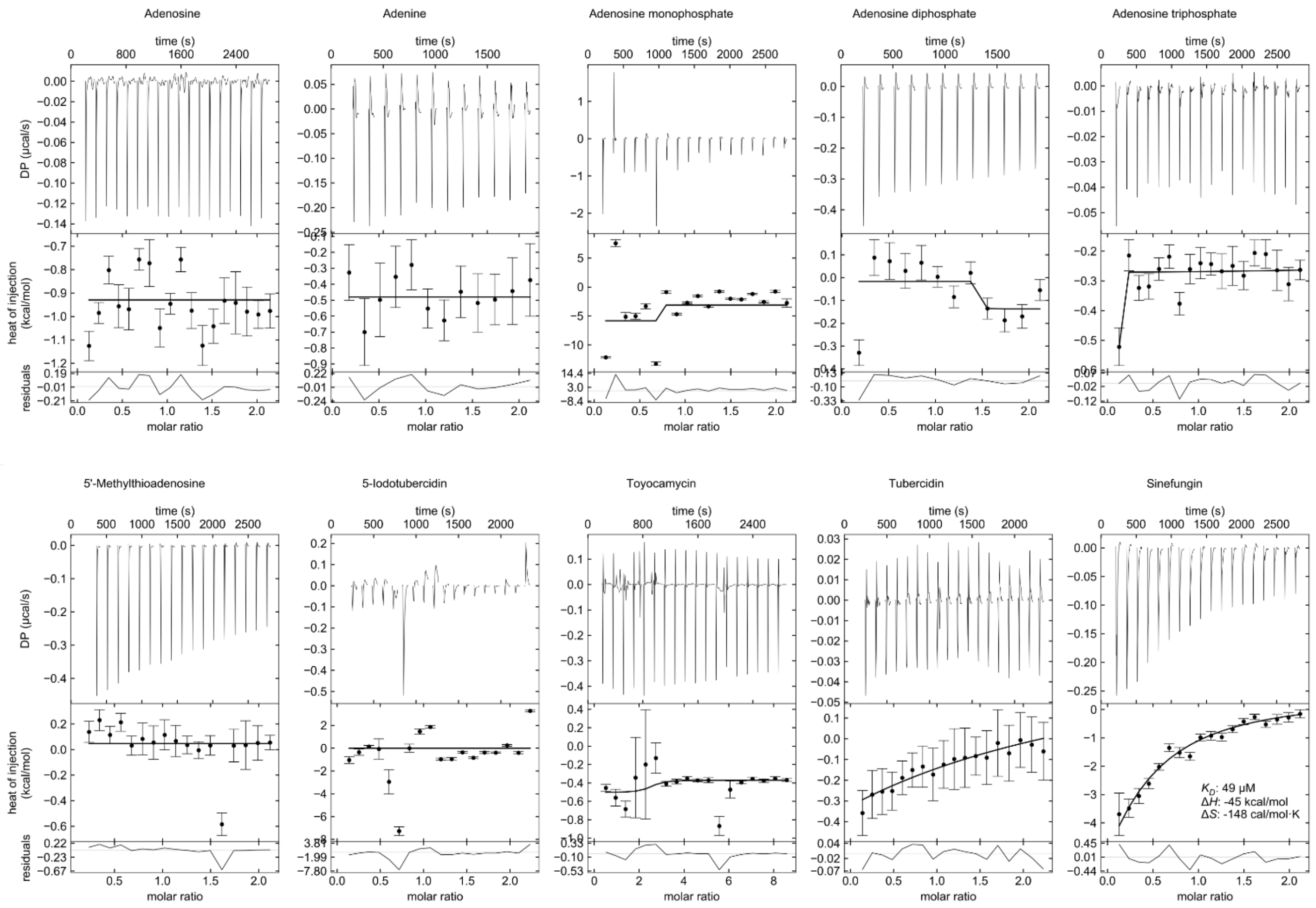


Figure S 10 **Affinities of compounds to nsp10-16 were measured by ITC with a protein concentration of 45 µM in the sample cell and compound concentration of 500 µM in the syringe.** All solutions contained the same buffer (50 mM Tris HCl, pH 7.5, 200 mM NaCl). Titrations of compound into buffer were performed as reference runs and subtracted from compound runs. Measurement of Toyocamycin was done with 2 mM compound concentration and all solutions were supplemented with 2% DMSO.

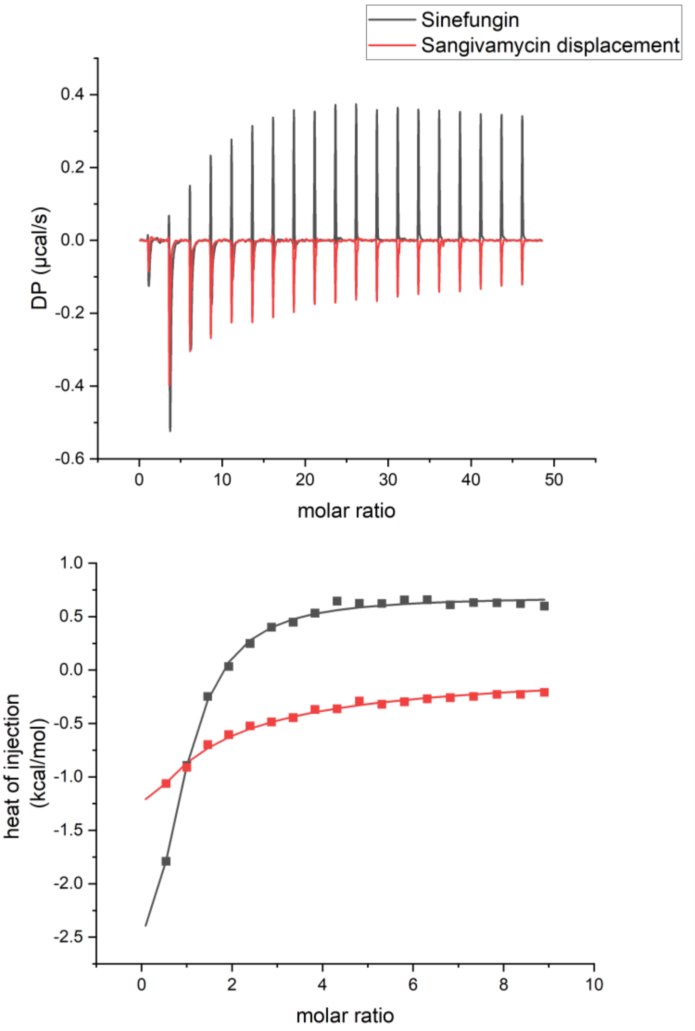


Figure S 11 **Overlay of ITC measurements for displacement of sangivamycin by sinefungin.** Sinefungin (2 mM) was titrated into 42.5 µM nsp10-16 mixed in advance with 2 mM sangivamycin. Buffer included 1 % DMSO. $K_D$ value for binding of sangivamycin was calculated with *ITCcalc* (Hammerschmidt et al. 2024) and is >1 mM. The measurement cannot give a more exact estimate of the $K_D$ value due to heavy precipitation after mixing of protein and sangivamycin which will have altered the protein and ligand concentrations in the mixture.

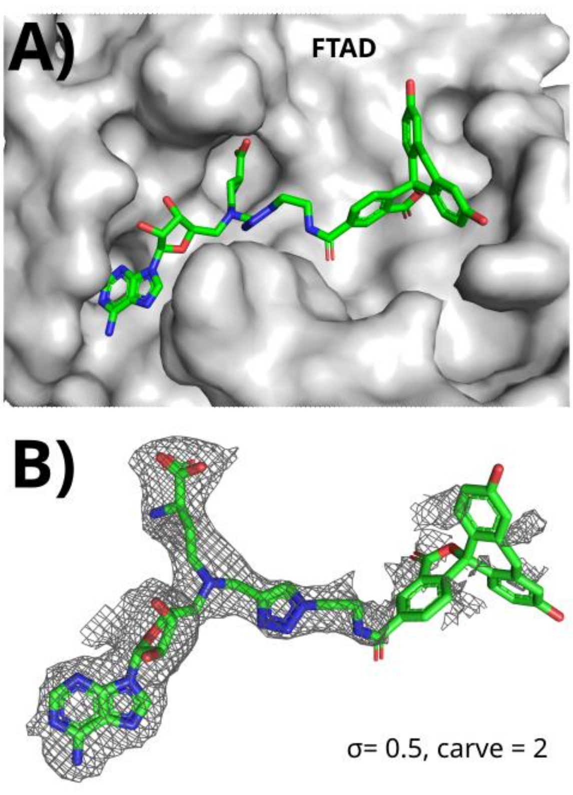


Figure S 12 **5-FAM-triazolyl-adenosyl-Dab (FTAD) bound to nsp10-16.** A) FTAD in stick representation and nsp16 SAM site as grey surface. B) Electron density of FTAD bound to nsp10-16.

#### Supplementary Tables

Table S1 Crystallographic table.

| PDB-ID | 8BSD | 8OV2 | 8OV3 | 8OV4 | 8BZV |
|---|---|---|---|---|---|
| **Compound** | **Tubercidin (TBN)** | **Sangivamycin (SGV)** | **5-Iodotubercidin (5ID)** | **Toyocamycin (TO1)** | **Adenosine (ADN)** |
| Wavelength (Å) | 1.033 | 1.033 | 1.033 | 1.033 | 1.033 |
| ***Processing statistics*** | | | | | |
| Resolution range (Å) | 37.54 - 1.95 (2.02 - 1.95) | 48.62 - 1.86 (1.93 - 1.86) | 48.21 - 1.82 (1.89 - 1.82) | 48.73 - 1.93 (2.00 - 1.93) | 42 - 1.8 (1.86 - 1.8) |
| Space group | $P\ 3_1\ 2\ 1$ | $P\ 3_1\ 2\ 1$ | $P\ 3_1\ 2\ 1$ | $P\ 3_1\ 2\ 1$ | $P\ 3_1\ 2\ 1$ |
| Unit cell parameters a, b, c (Å) | 167.47 167.47 51.51 | 168.02 168.02 51.58 | 167.01 167.01 51.4 | 168.05 168.05 51.71 | 167.32 167.32 51.54 |
| α, β, γ (°) | 90 90 120 | 90 90 120 | 90 90 120 | 90 90 120 | 90 90 120 |
| No. of reflections | 488472 (49468) | 569041 (53530) | 599801 (61005) | 510935 (51711) | 669170 (68246) |
| No. of unique reflections | 60413 (5982) | 70075 (6944) | 72530 (7171) | 62937 (6239) | 76612 (7609) |
| Multiplicity | 8.1 (8.3) | 8.1 (7.7) | 8.3 (8.5) | 8.1 (8.3) | 8.7 (9.0) |
| Completeness (%) | 99.87 (99.67) | 99.94 (100.00) | 98.44 (98.31) | 99.90 (100.00) | 99.96 (99.99) |
| mean $I/\sigma\ (I)$ | 8.68 (0.97) | 9.75 (0.84) | 7.33 (0.66) | 9.10 (0.80) | 9.71 (0.79) |
| Wilson B-factor ($Å^2$) | 37.25 | 35.57 | 32.08 | 38.46 | 29.92 |
| Rmerge (%) | 0.1756 (1.557) | 121.4 (1603) | 167.9 (2111) | 137.1 (1973) | 143.4 (1878) |
| Rmeas | 0.1874 (1.659) | 0.1296 (1.717) | 0.1791 (2.244) | 0.1465 (2.104) | 0.1526 (1.994) |
| Rp.i.m. | 0.0647 (0.5657) | 0.0449 (0.6079) | 0.0614 (0.7583) | 0.0509 (0.7243) | 0.0515 (0.6652) |
| $CC_{1/2}$ | 0.997 (0.465) | 0.998 (0.433) | 0.996 (0.4) | 0.997 (0.398) | 0.998 (0.395) |
| CC* | 0.999 (0.797) | 0.999 (0.777) | 0.999 (0.756) | 0.999 (0.755) | 0.999 (0.752) |
| ***Refinement statistics*** | | | | | |
| Reflections used in refinement | 60374 (5963) | 70066 (6944) | 72513 (7170) | 62932 (6239) | 76603 (7609) |
| Reflections used for $R_{free}$ | 634 (63) | 734 (71) | 741 (60) | 651 (55) | 804 (80) |
| $R_{work}$ | 0.1805 (0.3189) | 0.1773 (0.3307) | 0.1816 (0.3686) | 0.1791 (0.3328) | 0.1717 (0.3611) |
| $R_{free}$ | 0.2058 (0.3417) | 0.1995 (0.4185) | 0.2095 (0.3593) | 0.2075 (0.3586) | 0.1939 (0.3540) |
| *Number of atoms* | | | | | |
| Macromolecules | 3295 | 3277 | 3241 | 3268 | 3342 |
| Ligands | 405 | 284 | 270 | 281 | 482 |
| Solvent | 313 | 248 | 251 | 238 | 312 |
| Number of protein residues | 417 | 415 | 415 | 417 | 416 |
| RMS(bonds) | 0.011 | 0.011 | 0.011 | 0.012 | 0.011 |
| RMS(angles) | 1.22 | 1.13 | 1.16 | 1.2 | 1.23 |
| Ramachandran favored (%) | 96.61 | 97.57 | 97.08 | 96.85 | 97.82 |
| Ramachandran allowed (%) | 3.15 | 2.43 | 2.92 | 3.15 | 2.18 |
| Ramachandran outliers (%) | 0.24 | 0 | 0 | 0 | 0 |
| Rotamer outliers (%) | 0.83 | 0 | 0.56 | 0.56 | 0.82 |
| Clashscore | 2.89 | 4.29 | 3.45 | 3.71 | 2.68 |
| *Average B-factor ($Å^2$)* | | | | | |
| overall | 45.63 | 45.67 | 40.62 | 47.81 | 37.52 |
| Ligands | 59.55 | 55.95 | 49.24 | 58.22 | 51.69 |

| PDB-ID | 8OTO | 8OV1 | 8OSX | 8C5M | 8OT0 |
|---|---|---|---|---|---|
| **Compound** | **AMP** | **ADP** | **ATP** | **Methylthioadenosine (MTA)** | **MTA + Gly** |
| Wavelength (Å) | 0.976 | 0.976 | 1.033 | 0.976 | 0.976 |
| ***Processing statistics*** | | | | | |
| Resolution range (Å) | 42.07 - 1.8 (1.86 - 1.8) | 43.87 - 1.67 (1.73 - 1.67) | 42.14 - 1.83 (1.90 - 1.83) | 48.43 - 1.9 (1.97 - 1.9) | 43.77 - 2.21 (2.29 - 2.21) |
| Space group | $P\ 3_1\ 2\ 1$ | $P\ 3_1\ 2\ 1$ | $P\ 3_1\ 2\ 1$ | $P\ 3_1\ 2\ 1$ | $P\ 3_1\ 2\ 1$ |
| Unit cell parameters a, b, c (Å) | 167.86 167.86 51.592 | 167.495 167.495 51.5 | 168.02 168.02 51.7 | 166.955 166.955 51.394 | 167.102 167.102 51.379 |
| α, β, γ (°) | 90 90 120 | 90 90 120 | 90 90 120 | 90 90 120 | 90 90 120 |
| No. of reflections | 154355 (15306) | 191710 (18990) | 600082 (61240) | 1116167 (108977) | 82664 (8094) |
| No. of unique reflections | 77178 (7540) | 95855 (9407) | 73616 (7157) | 64765 (6253) | 41332 (4047) |
| Multiplicity | 2.0 (2.0) | 2.0 (2.0) | 8.2 (8.4) | 17.2 (17.0) | 2.0 (2.0) |
| Completeness (%) | 99.82 (98.51) | 99.87 (99.04) | 99.55 (97.77) | 99.63 (97.27) | 99.86 (99.14) |
| mean $I/\sigma\ (I)$ | 12.65 (0.76) | 12.57 (0.80) | 6.85 (0.27) | 13.34 (1.36) | 7.36 (0.89) |
| Wilson B-factor ($Å^2$) | 31.95 | 26.17 | 37.7 | 32.16 | 38.26 |
| Rmerge (%) | 33.5 (921.3) | 34.87 (847.1) | 195.1 (3118) | 153.1 (1755) | 76.41 (806.2) |
| Rmeas | 0.0473 (1.303) | 0.0493 (1.198) | 0.2083 (3.324) | 0.1578 (1.81) | 0.1081 (1.14) |
| Rp.i.m. | 0.0335 (0.9213) | 0.0349 (0.8471) | 0.0717 (1.138) | 0.0379 (0.4387) | 0.0764 (0.8062) |
| $CC_{1/2}$ | 0.999 (0.418) | 0.999 (0.5) | 0.996 (0.18) | 0.999 (0.645) | 0.995 (0.392) |
| CC* | 1 (0.768) | 1 (0.817) | 0.999 (0.552) | 1 (0.886) | 0.999 (0.75) |
| ***Refinement statistics*** | | | | | |
| Reflections used in refinement | 77046 (7539) | 95739 (9404) | 73444 (7156) | 64532 (6247) | 41326 (4046) |
| Reflections used for $R_{free}$ | 783 (73) | 988 (99) | 771 (83) | 690 (75) | 435 (44) |
| $R_{work}$ | 0.1755 (0.4075) | 0.1751 (0.3849) | 0.1973 (0.4356) | 0.1836 (0.3352) | 0.1863 (0.2990) |
| $R_{free}$ | 0.1857 (0.4028) | 0.1884 (0.3753) | 0.2221 (0.4271) | 0.2115 (0.3402) | 0.2163 (0.3175) |
| *Number of atoms* | | | | | |
| Macromolecules | 3333 | 3333 | 3336 | 3262 | 3255 |
| Ligands | 353 | 391 | 200 | 277 | 152 |
| Solvent | 275 | 329 | 238 | 272 | 190 |
| Number of protein residues | 417 | 417 | 417 | 416 | 417 |
| RMS(bonds) | 0.011 | 0.01 | 0.008 | 0.009 | 0.007 |
| RMS(angles) | 1.17 | 1.14 | 1.01 | 1.03 | 0.93 |
| Ramachandran favored (%) | 96.85 | 97.58 | 96.34 | 97.33 | 96.36 |
| Ramachandran allowed (%) | 2.91 | 2.42 | 3.66 | 2.67 | 3.64 |
| Ramachandran outliers (%) | 0.24 | 0 | 0 | 0 | 0 |
| Rotamer outliers (%) | 0 | 0 | 0.54 | 0 | 1.4 |
| Clashscore | 3.16 | 3.7 | 11.41 | 5.98 | 6.21 |
| *Average B-factor ($Å^2$)* | | | | | |
| overall | 38.56 | 32.74 | 42.72 | 36.2 | 43.34 |
| Ligands | 53.98 | 44.62 | 51.98 | 47.3 | 53.15 |

| PDB-ID | 8OTR | 9EMV | 8S8W | 9EMJ | 8S8X |
|---|---|---|---|---|---|
| **Compound** | **W08** | **SGV + Cap0-analog** | **SGV + Cap+-RNA** | **TO1 + Cap0-analog** | **TO1 + Cap0-RNA** |
| Wavelength (Å) | 1.033 | 0.827 | 1.033 | 1.033 | 1.033 |
| ***Processing statistics*** | | | | | |
| Resolution range (Å) | 41.99 - 1.77 (1.83 - 1.77) | 44.48 - 2.34 (2.42 - 2.34) | 48.8 - 2.10 (2.19 - 2.1) | 44.42 - 1.79 (1.85 - 1.79) | 49.06 - 1.994 (2.10 - 1.99) |
| Space group | $P\ 3_1\ 2\ 1$ | $P\ 3_1\ 2\ 1$ | $P\ 3_1\ 2\ 1$ | $P\ 3_1\ 2\ 1$ | $P\ 3_1\ 2\ 1$ |
| Unit cell parameters a, b, c (Å) | 167.6 167.6 51.48 | 169.51 169.51 52.25 | 167.14 167.14 51.83 | 168.7 168.7 52.25 | 168.11 168.11 52.11 |
| α, β, γ (°) | 90 90 120 | 90 90 120 | 90 90 120 | 90 90 120 | 90 90 120 |
| No. of reflections | 161333 (15956) | 848124 (82237) | 1123025 (140327) | 936249 (95368) | 1346340 (194729) |
| No. of unique reflections | 80684 (7847) | 36510 (3616) | 94499 (11830) | 79674 (7889) | 113051 (16215) |
| Multiplicity | 2.0 (2.0) | 23.2 (22.7) | 11.9 (11.9) | 11.8 (12.1) | 11.9 (12.0) |
| Completeness (%) | 99.79 (98.26) | 99.39 (97.10) | 99.78 (99.75) | 99.20 (98.38) | 97.47 (82.31) |
| mean $I/\sigma\ (I)$ | 8.41 (0.27) | 6.90 (0.48) | 6.88 (0.45) | 9.22 (0.53) | 4.64 (0.14) |
| Wilson B-factor ($Å^2$) | 35.9 | 49.01 | 45.5 | 31.54 | 45.88 |
| Rmerge (%) | 56.13 (2541) | 391.2 (3223) | 252.7 (3550) | 170.3 (2292) | 331.4 (3205) |
| Rmeas | 0.07937 (3.594) | 0.3998 (3.296) | 0.264 (3.708) | 0.178 (2.391) | 0.3462 (3.346) |
| Rp.i.m. | 0.0561 (2.541) | 0.0820 (0.6861) | 0.0760 (1.064) | 0.0514 (0.6777) | 0.0998 (0.9586) |
| $CC_{1/2}$ | 0.998 (0.158) | 0.997 (0.58) | 0.996 (0.227) | 0.998 (0.427) | 0.993 (0.271) |
| CC* | 1 (0.522) | 0.999 (0.857) | 0.999 (0.609) | 0.999 (0.773) | 0.998 (0.653) |
| ***Refinement statistics*** | | | | | |
| Reflections used in refinement | 80531 (7846) | 36304 (3515) | 48570 (6029) | 79603 (7835) | 56216 (6735) |
| Reflections used for $R_{free}$ | 825 (70) | 1809 (174) | 1080 (134) | 1034 (98) | 1047 (127) |
| $R_{work}$ | 0.1894 (0.4127) | 0.2029 (0.3805) | 0.1871 (0.3172) | 0.1810 (0.4539) | 0.1893 (0.4151) |
| $R_{free}$ | 0.2117 (0.3952) | 0.2567 (0.4051) | 0.2237 (0.3355) | 0.2070 (0.5026) | 0.2232 (0.3991) |
| *Number of atoms* | | | | | |
| Macromolecules | 3311 | 3263 | 3264 | 3269 | 3247 |
| Ligands | 193 | 303 | 144 | 208 | 112 |
| Solvent | 262 | 151 | 183 | 365 | 249 |
| Number of protein residues | 417 | 416 | 414 | 416 | 415 |
| RMS(bonds) | 0.107 | 0.013 | 0.008 | 0.011 | 0.008 |
| RMS(angles) | 1.66 | 1.28 | 0.95 | 1.09 | 0.98 |
| Ramachandran favored (%) | 96.85 | 96.35 | 95.84 | 98.06 | 96.59 |
| Ramachandran allowed (%) | 3.15 | 3.65 | 4.16 | 1.94 | 3.41 |
| Ramachandran outliers (%) | 0 | 0 | 0 | 0 | 0 |
| Rotamer outliers (%) | 0.27 | 0.28 | 0.56 | 0.28 | 0 |
| Clashscore | 8.81 | 5.83 | 3.13 | 1.96 | 1.51 |
| *Average B-factor ($Å^2$)* | | | | | |
| overall | 41.66 | 58.75 | 53.41 | 36.55 | 52.01 |
| Ligands | 49.62 | 61.84 | 71.31 | 40.22 | 56.01 |

| PDB-ID | 9EML |
|---|---|
| Compound | SAM + Cap0- analog (EDTA) |
| Wavelength (Å) | 1.033 |
| ***Processing statistics*** | |
| Resolution range (Å) | 49.07 - 2.4 (2.49 - 2.4) |
| Space group | *P* $3_1$ 2 1 |
| Unit cell parameters a, b, c (Å) | 167.74 167.74 52.13 |
| α, β, γ (°) | 90 90 120 |
| No. of reflections | 269215 (22747) |
| No. of unique reflections | 33107 (3298) |
| Multiplicity | 8.1 (6.9) |
| Completeness (%) | 99.90 (99.58) |
| mean $I/\sigma$ $(I)$ | 6.15 (0.66) |
| Wilson B-factor (Å$^2$) | 46.34 |
| Rmerge (%) | 315.7 (2637) |
| Rmeas | 0.3371 (2.851) |
| Rp.i.m. | 0.1167 (1.073) |
| $CC_{1/2}$ | 0.988 (0.232) |
| CC* | 0.997 (0.613) |
| ***Refinement statistics*** | |
| Reflections used in refinement | 33086 (3288) |
| Reflections used for $R_{free}$ | 527 (52) |
| $R_{work}$ | 0.1787 (0.2974) |
| $R_{free}$ | 0.2420 (0.3690) |
| *Number of atoms* | |
| Macromolecules | 3272 |
| Ligands | 397 |
| Solvent | 235 |
| Number of protein residues | 416 |
| RMS(bonds) | 0.012 |
| RMS(angles) | 1.23 |
| Ramachandran favored (%) | 96.09 |
| Ramachandran allowed (%) | 3.91 |
| Ramachandran outliers (%) | 0 |
| Rotamer outliers (%) | 0.28 |
| Clashscore | 3.08 |
| *Average B-factor (Å$^2$)* | |
| overall | 53.42 |
| Ligands | 64.44 |

Table S2 List of screened compounds
*Separate file*

Table S3. Matches of a SiteMine similarity search with the ligand binding site of WZ16 in PDB entry 7r1u as query. Matches up to a score of 31 are shown.

| PDB Entry | SP-Score | Pharmacophore Score | Shape Score | Ligandname | Similarity Score | Protein Name | UniProt Accession | Cluster |
|---|---|---|---|---|---|---|---|---|
| 2NYU | 51.6 | 24.6 | 27 | SAM_B_201 | 47.0 | rRNA methyltransferase 2, mitochondrial | Q9UI43 | 1 |
| 2NYU | 50.6 | 23.6 | 27 | SAM_A_201 | 44.2 | rRNA methyltransferase 2, mitochondrial | Q9UI43 | 1 |
| 7O9K | 45.4 | 20.4 | 25 | SAH_n_301 | 37.0 | rRNA methyltransferase 2, mitochondrial | Q9UI43 | 1 |
| 4N49 | 50.6 | 23.6 | 27 | SAM_A_601 | 44.2 | Cap-specific mRNA (nucleoside-2'-O-)-methyltransferase 1 | Q8N1G2 | 2 |
| 4N48 | 50.6 | 22.6 | 28 | SAM_B_601 | 40.8 | Cap-specific mRNA (nucleoside-2'-O-)-methyltransferase 1 | Q8N1G2 | 2 |
| 4N48 | 47.2 | 21.2 | 26 | SAM_A_601 | 38.5 | Cap-specific mRNA (nucleoside-2'-O-)-methyltransferase 1 | Q8N1G2 | 2 |
| 8P4F | 42.8 | 18.8 | 24 | SAM_O_901 | 33.5 | Cap-specific mRNA (nucleoside-2'-O-)-methyltransferase 1 | Q8N1G2 | 2 |
| 7YRI | 45.8 | 21.8 | 24 | SAM_C_401 | 41.6 | S-adenosylmethionine sensor upstream of mTORC1 | | 3 |
| 7YRI | 46.8 | 21.8 | 25 | SAM_D_401 | 40.8 | S-adenosylmethionine sensor upstream of mTORC1 | | 3 |
| 7YRI | 44.8 | 19.8 | 25 | SAM_A_401 | 35.5 | S-adenosylmethionine sensor upstream of mTORC1 | | 3 |
| 7YRI | 43.4 | 19.4 | 24 | SAM_J_401 | 35.1 | S-adenosylmethionine sensor upstream of mTORC1 | | 3 |
| 3P71 | 47.8 | 21.8 | 26 | AN6_C_310 | 40.1 | Leucine carboxyl methyltransferase 1 | Q9UIC8 | 4 |
| 3IEI | 48.6 | 21.6 | 27 | SAH_A_601 | 38.9 | Leucine carboxyl methyltransferase 1 | Q9UIC8 | 4 |
| 3IEI | 46.8 | 20.8 | 26 | SAH_H_601 | 37.4 | Leucine carboxyl methyltransferase 1 | Q9UIC8 | 4 |
| 3O7W | 46.4 | 20.4 | 26 | SAM_A_801 | 36.4 | Leucine carboxyl methyltransferase 1 | Q9UIC8 | 4 |
| 3IEI | 44.8 | 19.8 | 25 | SAH_D_601 | 35.5 | Leucine carboxyl methyltransferase 1 | Q9UIC8 | 4 |
| 3IEI | 44.8 | 19.8 | 25 | SAH_G_601 | 35.5 | Leucine carboxyl methyltransferase 1 | Q9UIC8 | 4 |
| 3IEI | 44.8 | 19.8 | 25 | SAH_F_601 | 35.5 | Leucine carboxyl methyltransferase 1 | Q9UIC8 | 4 |
| 3IEI | 45.8 | 19.8 | 26 | SAH_B_601 | 34.9 | Leucine carboxyl methyltransferase 1 | Q9UIC8 | 4 |
| 3IEI | 43.8 | 18.8 | 25 | SAH_C_601 | 32.9 | Leucine carboxyl methyltransferase 1 | Q9UIC8 | 4 |
| 3GDH | 41.2 | 20.2 | 21 | MGP_C_3 | 39.6 | Trimethylguanosine synthase TGS1 | Q96RS0 | 5 |
| 3EGI | 43.2 | 20.2 | 23 | ADP_C_1 | 37.9 | Trimethylguanosine synthase TGS1 | Q96RS0 | 5 |
| 3EGI | 41.2 | 19.2 | 22 | ADP_D_2 | 36.0 | Trimethylguanosine synthase TGS1 | Q96RS0 | 5 |
| 3GDH | 37.2 | 17.2 | 20 | MGP_B_2 | 32.0 | Trimethylguanosine synthase TGS1 | Q96RS0 | 5 |

| | | | | | | | | |
|---|---|---|---|---|---|---|---|---|
| 3EGI | 39.4 | 17.4 | 22 | ADP_A_3 | 31.2 | Trimethylguanosine synthase TGS1 | Q96RS0 | 5 |
| 5WWQ | 43.8 | 20.8 | 23 | | 39.6 | tRNA (cytosine(72)-C(5))-methyltransferase NSUN6 | Q8TEA1 | 6 |
| 5WWT | 42.8 | 19.8 | 23 | | 36.8 | tRNA (cytosine(72)-C(5))-methyltransferase NSUN6 | Q8TEA1 | 6 |
| 2B9E | 40 | 19 | 21 | SAM_A_1201 | 36.2 | tRNA (cytosine(72)-C(5))-methyltransferase NSUN6 | Q8TEA1 | 6 |
| 5WWS | 39.8 | 18.8 | 21 | SAM_A_501 | 35.6 | tRNA (cytosine(72)-C(5))-methyltransferase NSUN6 | Q8TEA1 | 6 |
| 5WWS | 39.8 | 18.8 | 21 | SAM_B_501 | 35.6 | tRNA (cytosine(72)-C(5))-methyltransferase NSUN6 | Q8TEA1 | 6 |
| 5WWT | 38 | 18 | 20 | | 34.2 | tRNA (cytosine(72)-C(5))-methyltransferase NSUN6 | Q8TEA1 | 6 |
| 5WWQ | 39 | 18 | 21 | | 33.4 | tRNA (cytosine(72)-C(5))-methyltransferase NSUN6 | Q8TEA1 | 6 |
| 5C9Z | 40.8 | 19.8 | 21 | SFG_A_701 | 38.5 | Protein arginine N-methyltransferase 5 | O14744 | 7 |
| 5EMM | 40.8 | 19.8 | 21 | SFG_A_701 | 38.5 | Protein arginine N-methyltransferase 5 | O14744 | 7 |
| 6CKC | 37.8 | 18.8 | 19 | F5J_A_701 | 37.4 | Protein arginine N-methyltransferase 5 | O14744 | 7 |
| 7ZUY | 37.8 | 18.8 | 19 | MTA_A_701 | 37.4 | Protein arginine N-methyltransferase 5 | O14744 | 7 |
| 7ZVU | 37.8 | 18.8 | 19 | MTA_A_701 | 37.4 | Protein arginine N-methyltransferase 5 | O14744 | 7 |
| 4X61 | 38.8 | 18.8 | 20 | SAM_A_701 | 36.5 | Protein arginine N-methyltransferase 5 | O14744 | 7 |
| 6K1S | 38.8 | 18.8 | 20 | CUX_A_700 | 36.5 | Protein arginine N-methyltransferase 5 | O14744 | 7 |
| 7ZUQ | 38.8 | 18.8 | 20 | MTA_A_701 | 36.5 | Protein arginine N-methyltransferase 5 | O14744 | 7 |
| 7ZUU | 40 | 19 | 21 | MTA_A_701 | 36.2 | Protein arginine N-methyltransferase 5 | O14744 | 7 |
| 6V0O | 36 | 18 | 18 | SFG_A_701 | 36.0 | Protein arginine N-methyltransferase 5 | O14744 | 7 |
| 6V0P | 39.8 | 18.8 | 21 | SFG_A_704 | 35.6 | Protein arginine N-methyltransferase 5 | O14744 | 7 |
| 4GQB | 35.8 | 17.8 | 18 | 0XU_A_701 | 35.4 | Protein arginine N-methyltransferase 5 | O14744 | 7 |
| 7ZUP | 35.8 | 17.8 | 18 | MTA_A_701 | 35.4 | Protein arginine N-methyltransferase 5 | O14744 | 7 |
| 4X63 | 37 | 18 | 19 | SAH_A_701 | 35.1 | Protein arginine N-methyltransferase 5 | O14744 | 7 |
| 6RLL | 37.8 | 17.8 | 20 | K8H_A_701 | 33.6 | Protein arginine N-methyltransferase 5 | O14744 | 7 |
| 6V0N | 37.8 | 17.8 | 20 | SFG_A_701 | 33.6 | Protein arginine N-methyltransferase 5 | O14744 | 7 |
| 7KIC | 37.8 | 17.8 | 20 | WFS_A_706 | 33.6 | Protein arginine N-methyltransferase 5 | O14744 | 7 |
| 8G1U | 37.8 | 17.8 | 20 | ADN_E_701 | 33.6 | Protein arginine N-methyltransferase 5 | O14744 | 7 |

| | | | | | | | | |
|---|---|---|---|---|---|---|---|---|
| 5EMK | 38.8 | 17.8 | 21 | SFG_A_701 | 32.9 | Protein arginine N-methyltransferase 5 | O14744 | 7 |
| 7KID | 34.8 | 16.8 | 18 | WFV_A_705 | 32.5 | Protein arginine N-methyltransferase 5 | O14744 | 7 |
| 7MXA | 34.8 | 16.8 | 18 | ZR4_A_701 | 32.5 | Protein arginine N-methyltransferase 5 | O14744 | 7 |
| 7SES | 34.8 | 16.8 | 18 | 97X_A_701 | 32.5 | Protein arginine N-methyltransferase 5 | O14744 | 7 |
| 8G1U | 34.8 | 16.8 | 18 | ADN_A_701 | 32.5 | Protein arginine N-methyltransferase 5 | O14744 | 7 |
| 7UY1 | 35.8 | 16.8 | 19 | PJ0_A_701 | 31.7 | Protein arginine N-methyltransferase 5 | O14744 | 7 |
| 6RLQ | 31.8 | 15.8 | 16 | K8N_A_701 | 31.4 | Protein arginine N-methyltransferase 5 | O14744 | 7 |
| 5E9W | 48.4 | 21.4 | 27 | SAH_C_501 | 38.4 | mRNA cap guanine-N7 methyltransferase | O43148 | 8 |
| 3BGV | 43.2 | 20.2 | 23 | SAH_C_313 | 37.9 | mRNA cap guanine-N7 methyltransferase | O43148 | 8 |
| 5E9J | 43.2 | 20.2 | 23 | SAH_B_501 | 37.9 | mRNA cap guanine-N7 methyltransferase | O43148 | 8 |
| 5E9J | 40.4 | 19.4 | 21 | SAH_A_501 | 37.3 | mRNA cap guanine-N7 methyltransferase | O43148 | 8 |
| 3BGV | 43.4 | 19.4 | 24 | SAH_C_315 | 35.1 | mRNA cap guanine-N7 methyltransferase | O43148 | 8 |
| 5E9W | 40.6 | 18.6 | 22 | SAH_D_501 | 34.3 | mRNA cap guanine-N7 methyltransferase | O43148 | 8 |
| 5E8J | 37.6 | 17.6 | 20 | GOL_A_501 | 33.1 | mRNA cap guanine-N7 methyltransferase | O43148 | 8 |
| 5E8J | 37.6 | 17.6 | 20 | SAH_B_501 | 33.1 | mRNA cap guanine-N7 methyltransferase | O43148 | 8 |
| 3BGV | 41.2 | 18.2 | 23 | SAH_C_314 | 32.6 | mRNA cap guanine-N7 methyltransferase | O43148 | 8 |
| 5E9W | 42.2 | 18.2 | 24 | SAH_B_501 | 32.0 | mRNA cap guanine-N7 methyltransferase | O43148 | 8 |
| 5E9W | 42.2 | 18.2 | 24 | SAH_A_500 | 32.0 | mRNA cap guanine-N7 methyltransferase | O43148 | 8 |
| 3BGV | 39.6 | 17.6 | 22 | SAH_C_316 | 31.7 | mRNA cap guanine-N7 methyltransferase | O43148 | 8 |
| 6DCB | 40.6 | 19.6 | 21 | SAH_A_701 | 37.9 | 7SK snRNA methylphosphate capping enzyme | Q7L2J0 | 9 |
| 5UNA | 41.6 | 19.6 | 22 | SAH_A_701 | 37.1 | 7SK snRNA methylphosphate capping enzyme | Q7L2J0 | 9 |
| 5UNA | 38.6 | 18.6 | 20 | SAH_F_701 | 35.9 | 7SK snRNA methylphosphate capping enzyme | Q7L2J0 | 9 |
| 5UNA | 36.6 | 17.6 | 19 | SAH_B_701 | 33.9 | 7SK snRNA methylphosphate capping enzyme | Q7L2J0 | 9 |
| 6DCC | 37.6 | 17.6 | 20 | SAH_A_701 | 33.1 | 7SK snRNA methylphosphate capping enzyme | Q7L2J0 | 9 |

| | | | | | | | | |
|---|---|---|---|---|---|---|---|---|
| 5UNA | 35.6 | 16.6 | 19 | SAH_C_701 | 31.1 | 7SK snRNA methylphosphate capping enzyme | Q7L2J0 | 9 |
| 6KHS | 41.8 | 19.8 | 22 | MEQ_A_301 | 37.6 | Methyltransferase N6AMT1 | Q9Y5N5 | 10 |
| 6KMS | 40.6 | 18.6 | 22 | SAM_C_301 | 34.3 | Methyltransferase N6AMT1 | Q9Y5N5 | 10 |
| 6H1D | 41.6 | 18.6 | 23 | SAH_A_301 | 33.6 | Methyltransferase N6AMT1 | Q9Y5N5 | 10 |
| 6KMR | 38.8 | 17.8 | 21 | SAM_B_301 | 32.9 | Methyltransferase N6AMT1 | Q9Y5N5 | 10 |
| 1U7T | 44 | 20 | 24 | TDT_A_501 | 36.7 | 3-hydroxyacyl-CoA dehydrogenase type-2 | Q99714 | 11 |
| 3QOW | 44 | 20 | 24 | SAM_A_417 | 36.7 | Histone-lysine N-methyltransferase, H3 lysine-79 specific | Q8TEK3 | 12 |
| 1NW3 | 42.4 | 19.4 | 23 | SAM_A_500 | 35.8 | Histone-lysine N-methyltransferase, H3 lysine-79 specific | Q8TEK3 | 12 |
| 4U7T | 41.4 | 19.4 | 22 | SAH_A_1004 | 36.5 | DNA (cytosine-5)-methyltransferase 3A | Q9Y6K1 | 13 |
| 6F57 | 41.4 | 19.4 | 22 | SAH_D_1001 | 36.5 | DNA (cytosine-5)-methyltransferase 3A | Q9Y6K1 | 13 |
| 2QRV | 38.4 | 18.4 | 20 | SAH_E_5 | 35.3 | DNA (cytosine-5)-methyltransferase 3A | Q9Y6K1 | 13 |
| 6W89 | 38.4 | 18.4 | 20 | SAH_D_1002 | 35.3 | DNA (cytosine-5)-methyltransferase 3A | Q9Y6K1 | 13 |
| 6F57 | 39.4 | 18.4 | 21 | SAH_A_1001 | 34.5 | DNA (cytosine-5)-methyltransferase 3A | Q9Y6K1 | 13 |
| 2QRV | 39.2 | 18.2 | 21 | SAH_H_8 | 34.0 | DNA (cytosine-5)-methyltransferase 3A | Q9Y6K1 | 13 |
| 6W8J | 40.2 | 18.2 | 22 | SAH_A_1001 | 33.3 | DNA (cytosine-5)-methyltransferase 3A | Q9Y6K1 | 13 |
| 6W8D | 41.2 | 18.2 | 23 | SAH_A_1001 | 32.6 | DNA (cytosine-5)-methyltransferase 3A | Q9Y6K1 | 13 |
| 4U7T | 38.6 | 17.6 | 21 | SAH_C_1004 | 32.4 | DNA (cytosine-5)-methyltransferase 3A | Q9Y6K1 | 13 |
| 2QRV | 37.2 | 17.2 | 20 | SAH_A_1 | 32.0 | DNA (cytosine-5)-methyltransferase 3A | Q9Y6K1 | 13 |
| 6W8B | 39.4 | 17.4 | 22 | SAH_A_1001 | 31.2 | DNA (cytosine-5)-methyltransferase 3A | Q9Y6K1 | 13 |
| 6W8B | 39.4 | 17.4 | 22 | SAH_H_1001 | 31.2 | DNA (cytosine-5)-methyltransferase 3A | Q9Y6K1 | 13 |
| 6W8B | 39.4 | 17.4 | 22 | SAH_K_1001 | 31.2 | DNA (cytosine-5)-methyltransferase 3A | Q9Y6K1 | 13 |
| 6KDP | 41.4 | 19.4 | 22 | SAH_A_902 | 36.5 | DNA (cytosine-5)-methyltransferase 3B | Q9UBC3 | 14 |
| 7X9D | 42.2 | 19.2 | 23 | HRM_A_901 | 35.2 | DNA (cytosine-5)-methyltransferase 3B | Q9UBC3 | 14 |

| | | | | | | | | |
|---|---|---|---|---|---|---|---|---|
| 6U8W | 37.4 | 17.4 | 20 | SAH_D_901 | 32.5 | DNA (cytosine-5)-methyltransferase 3B | Q9UBC3 | 14 |
| 6KDT | 38.2 | 17.2 | 21 | SAH_D_901 | 31.3 | DNA (cytosine-5)-methyltransferase 3B | Q9UBC3 | 14 |
| 7PNZ | 42.4 | 19.4 | 23 | SAH_b_501 | 35.8 | 12S rRNA N4-methylcytidine (m4C) methyltransferase | A6NJ78 | 15 |
| 7PNY | 43.4 | 19.4 | 24 | SAH_b_501 | 35.1 | 12S rRNA N4-methylcytidine (m4C) methyltransferase | A6NJ78 | 15 |
| 7SFD | 42.4 | 19.4 | 23 | SAH_A_1701 | 35.8 | DNA (cytosine-5)-methyltransferase 1 | P26358 | 16 |
| 3SWR | 38.4 | 18.4 | 20 | SFG_A_300 | 35.3 | DNA (cytosine-5)-methyltransferase 1 | P26358 | 16 |
| 6X9I | 40.2 | 18.2 | 22 | SAH_A_1701 | 33.3 | DNA (cytosine-5)-methyltransferase 1 | P26358 | 16 |
| 7XI9 | 37.6 | 17.6 | 20 | SAH_A_1705 | 33.1 | DNA (cytosine-5)-methyltransferase 1 | P26358 | 16 |
| 4WXX | 39.4 | 17.4 | 22 | SAH_A_1706 | 31.2 | DNA (cytosine-5)-methyltransferase 1 | P26358 | 16 |
| 1ZQ9 | 41 | 19 | 22 | SAM_B_4001 | 35.4 | Probable dimethyladenosine transferase | Q9UNQ2 | 17 |
| 6L8U | 41 | 19 | 22 | SAH_C_1001 | 35.4 | RNA 5'-monophosphate methyltransferase | Q7Z5W3 | 18 |
| 6L8U | 42 | 19 | 23 | SAH_A_1001 | 34.7 | RNA 5'-monophosphate methyltransferase | Q7Z5W3 | 18 |
| 6L8U | 39.4 | 18.4 | 21 | SAH_B_1001 | 34.5 | RNA 5'-monophosphate methyltransferase | Q7Z5W3 | 18 |
| 6L8U | 42.6 | 18.6 | 24 | SAH_D_1001 | 33.0 | RNA 5'-monophosphate methyltransferase | Q7Z5W3 | 18 |
| 7EU5 | 43.4 | 19.4 | 24 | SAH_C_301 | 35.1 | Nicotinamide N-methyltransferase | P40261 | 19 |
| 7ET7 | 38.2 | 17.2 | 21 | SAH_B_301 | 31.3 | Nicotinamide N-methyltransferase | P40261 | 19 |
| 5CCB | 39.6 | 18.6 | 21 | SAH_A_301 | 35.1 | tRNA (adenine(58)-N(1))-methyltransferase catalytic subunit TRMT61A | Q96FX7 | 20 |
| 7WTU | 35.6 | 17.6 | 18 | SAH_q_301 | 34.8 | Small ribosomal subunit protein uS9 | P62249 | 21 |
| 7WTT | 40.2 | 18.2 | 22 | SAH_q_301 | 33.3 | Small ribosomal subunit protein uS9 | P62249 | 21 |
| 7WTV | 38.2 | 17.2 | 21 | SAH_q_301 | 31.3 | Small ribosomal subunit protein uS9 | P62249 | 21 |
| 3ORH | 44.4 | 19.4 | 25 | SAH_D_4003 | 34.5 | Guanidinoacetate N-methyltransferase | Q14353 | 22 |
| 3ORH | 37.6 | 17.6 | 20 | SAH_C_4002 | 33.1 | Guanidinoacetate N-methyltransferase | Q14353 | 22 |
| 3ORH | 37.6 | 17.6 | 20 | SAH_A_4000 | 33.1 | Guanidinoacetate N-methyltransferase | Q14353 | 22 |
| 7EZG | 44.2 | 19.2 | 25 | SAH_A_301 | 33.9 | tRNA N(3)-methylcytidine methyltransferase METTL6 | Q8TCB7 | 23 |
| 7F1E | 37 | 17 | 20 | SAM_B_301 | 31.5 | tRNA N(3)-methylcytidine | Q8TCB7 | 23 |

| | | | | | | | | |
|---|---|---|---|---|---|---|---|---|
| | | | | | | methyltransferase METTL6 | | |
| 3KQO | 39 | 18 | 21 | SAH_A_2001 | 33.4 | Phenylethanolamine N-methyltransferase | P11086 | 24 |
| 3KQW | 39 | 18 | 21 | SAH_A_2001 | 33.4 | Phenylethanolamine N-methyltransferase | P11086 | 24 |
| 3HCA | 36 | 17 | 19 | SAH_A_3001 | 32.2 | Phenylethanolamine N-methyltransferase | P11086 | 24 |
| 3HCB | 37 | 17 | 20 | SAH_A_2001 | 31.5 | Phenylethanolamine N-methyltransferase | P11086 | 24 |
| 3KPJ | 37 | 17 | 20 | SAH_A_3001 | 31.5 | Phenylethanolamine N-methyltransferase | P11086 | 24 |
| 3KQV | 37 | 17 | 20 | SAH_A_2001 | 31.5 | Phenylethanolamine N-methyltransferase | P11086 | 24 |
| 5E1O | 37.4 | 17.4 | 20 | SAH_B_301 | 32.5 | N-terminal Xaa-Pro-Lys N-methyltransferase 1 | Q9BV86 | 25 |
| 2B25 | 36 | 17 | 19 | SAM_A_601 | 32.2 | tRNA (adenine(58)-N(1))-methyltransferase, mitochondrial | Q9BVS5 | 26 |
| 7K3D | 34.6 | 16.6 | 18 | SAH_B_301 | 31.9 | tRNA (adenine(58)-N(1))-methyltransferase, mitochondrial | Q9BVS5 | 26 |
| 5U4X | 38.4 | 17.4 | 21 | SAH_C_501 | 31.8 | Histone-arginine methyltransferase CARM1 | Q86X55 | 27 |
| 6AAX | 40.8 | 17.8 | 23 | SAM_A_401 | 31.6 | Dimethyladenosine transferase 1, mitochondrial | Q8WVM0 | 28 |
| 7SE7 | 42 | 18 | 24 | SAM_A_401 | 31.5 | rRNA 2'-O-methyltransferase fibrillarin | P22087 | 29 |
| 7SE8 | 41.8 | 17.8 | 24 | 8W1_B_401 | 31.0 | rRNA 2'-O-methyltransferase fibrillarin | P22087 | 29 |
| 4FP9 | 37 | 17 | 20 | SAM_C_401 | 31.5 | 5-methylcytosine rRNA methyltransferase NSUN4 | Q96CB9 | 30 |
| 6H2V | 35.6 | 16.6 | 19 | SAM_A_301 | 31.1 | rRNA N6-adenosine-methyltransferase METTL5 | Q9NRN9 | 31 |
| 8D58 | 35.6 | 16.6 | 19 | | 31.1 | tRNA (guanine-N(7)-)-methyltransferase | Q9UBP6 | 32 |
| 4FT2 | 38.8 | 17.8 | 21 | SAH_A_1000 | 32.9 | DNMT! (but plant) | Q9AXT8 | |
